# Motility Control Through an Anti-Activation Mechanism in *Agrobacterium tumefaciens*

**DOI:** 10.1101/2021.06.24.449765

**Authors:** Melene A. Alakavuklar, Brynn C. Heckel, Ari M. Stoner, Joseph A. Stembel, Clay Fuqua

## Abstract

Many bacteria can migrate from a free-living, planktonic state to an attached, biofilm existence. One factor regulating this transition in the facultative plant pathogen *Agrobacterium tumefaciens* is the ExoR-ChvG-ChvI system. Periplasmic ExoR regulates activity of the ChvG-ChvI two-component system in response to environmental stress, most notably low pH. ChvI impacts hundreds of genes, including those required for type VI secretion, virulence, biofilm formation, and flagellar motility. Previous studies revealed that activated ChvG-ChvI represses expression of most of class II and class III flagellar biogenesis genes, but not the master motility regulators *visN*, *visR*, and *rem*. In this study, we characterized the integration of the ExoR-ChvG-ChvI and VisNR-Rem pathways. We isolated motile suppressors of the non-motile Δ*exoR* mutant and thereby identified the previously unannotated *mirA* gene encoding a 76 amino acid protein. We report that the MirA protein interacts directly with the Rem DNA-binding domain, sequestering Rem and preventing motility gene activation. The ChvG-ChvI pathway activates *mirA* expression and elevated *mirA* is sufficient to block motility. This study reveals how the ExoR-ChvG-ChvI pathway prevents flagellar motility in *A. tumefaciens*. MirA is also conserved among other members of the *Rhizobiales* suggesting similar mechanisms of motility regulation.

**Plain language summary:** *Agrobacterium tumefaciens* is a plant pathogen that responds to low pH, stressful environments via a complex two-component regulatory system. This regulatory system can completely inhibit bacterial locomotion by flagella, and in this study we define the mechanism for this inhibition and identify a novel small protein regulator that blocks flagellar gene expression.

## Introduction

The bacterial flagellum is one of the most complex and sophisticated externalized structures that bacteria produce. Flagellar biosynthesis genes in bacteria are under hierarchical regulation, resulting in morphogenetic control of flagellum assembly (Chevance & Hughes, 2008, Soutourina & Bertin, 2003). The proper temporal and stoichiometric control of flagellar biogenesis dictates the inside-out assembly of flagella and provides checkpoints to ensure that the requisite flagellar components are produced in sufficient quantities as they are required (Chevance & Hughes, 2017). Based on the best-studied systems in *Escherichia coli* and *Salmonella enterica* as models, the top genes of the hierarchy (class I) encode transcription factors that activate expression of genes specifying the flagellar machinery components (class II) (Kawagishi *et al*., 1992). Class II genes encode proteins that make up the hook-basal body complex of the flagellum. The basal body includes the type III secretion system that exports hook and filament proteins outside of the cell for filament assembly and the torque-generating motor proteins required for filament rotation. Assembly of the basal body permits class II proteins to activate expression of class III genes (Chevance & Hughes, 2008). In the *Gammaproteobacteria* (such as *E. coli, S. enterica,* and *Pseudomonas aeruginosa*) and the Gram-positive bacterium *Bacillus subtilis*, class II genes encode the entire hook-basal body complex, and class III genes include hook-filament junction proteins and flagellin proteins (Mukherjee & Kearns, 2014).

In *A. tumefaciens* and other closely related *Alphaproteobacteria* (APB) in the order *Rhizobiales*, there are two tiers of class I transcription factors that initiate flagellar gene expression; these have been separated into class IA and class IB. Class IA consists of the LuxR-FixJ-type transcription factors VisN and VisR (Rotter *et al*., 2006, Sourjik *et al*., 2000, Tambalo *et al*., 2010, Xu *et al*., 2013). When either *visN* (ATU_RS02580) or *visR* (ATU_RS02585) are absent, transcription of flagellar biosynthesis and chemotaxis genes is abolished and cells are non-motile. VisN and VisR are predicted to function as heteromultimers because of the redundancy of their single mutant phenotypes (Sourjik *et al*., 2000). In *Sinorhizobium meliloti,* the motility gene expression defect of *visNR* mutants is directly caused by loss of expression of the class IB gene, *rem* (Rotter *et al*., 2006). Rem is an orphan OmpR-like transcription factor that is required for expression of all motility and chemotaxis genes. Rem from *S. meliloti* directly binds to and activates expression of several promoters that control flagellar basal body protein-encoding genes (Rotter *et al*., 2006). The hierarchical system of motility gene transcription in APB requires the class IB gene *rem* for expression of all class II and III flagellar and chemotaxis genes. Ectopic expression of *rem* in class IA *visNR* mutants is epistatic to the Δ*visNR* motility defect, driving normal motility and demonstrating the regulatory hierarchy in this pathway.

Flagellar assembly and swimming motility in *A. tumefaciens* and related rhizobia are repressed in acidic conditions via the pH-responsive ExoR-ChvG-ChvI signaling system (Heckel *et al*., 2014, Yao *et al*., 2004, Yuan *et al*., 2008). In neutral conditions, the periplasmic protein ExoR binds to the sensor kinase ChvG and prevents initiation of phosphotransfer to ChvI in this two-component system (Chen *et al*., 2008, Wells *et al*., 2007). Upon cultivation in low pH growth media, ExoR is proteolytically inactivated, unfettering the ChvGI system, and impacting a large regulon (Lu *et al*., 2012). In *A. tumefaciens,* the activity of ChvG-ChvI leads to dramatic decreases in motility gene expression (Heckel *et al*., 2014). The expression of nearly every motility-related gene is decreased when *A. tumefaciens* cells encounter acidic conditions, including both flagellar biosynthetic gene clusters and chemotaxis genes found throughout the *A. tumefaciens* genome. This phenomenon occurs whether cells are exposed to low pH media or in an Δ*exoR* (ATU_RS08400) mutant (Heckel *et al*., 2014, Yuan *et al*., 2008). The mechanism by which activated ChvG-ChvI so effectively represses motility gene expression has remained unclear. We hypothesize that the low pH-activated ChvG-ChvI system functions to disrupt the flagellar gene expression hierarchy.

Another complex phenotype that is regulated by environmental pH in *A. tumefaciens* is attachment to surfaces. The acid-responsive ExoR-ChvG-ChvI system has a dramatic effect on attachment and biofilm formation: Δ*exoR* mutants are completely deficient for attachment and fail to form biofilms on both abiotic and plant surfaces (Tomlinson *et al*., 2010). *A. tumefaciens* cells grown in acidic conditions also fail to form biofilms. The attachment defect of acid-exposed cells and Δ*exoR* mutants is dependent on the two-component system ChvG-ChvI, and mutation of either regulator gene in an Δ*exoR* mutant background restores biofilm formation capabilities (Heckel *et al*., 2014). Thus, an active ChvG-ChvI system serves to inhibit both swimming motility and biofilm formation. The uniformly negative impact that active ChvG-ChvI has on motility and attachment is atypical, because these two complex phenotypes are often considered as inverse processes; that is, bacteria will progress through either a motile or sessile program depending on the chemical profile of their environment (Prüß, 2017). The VisN and VisR regulators have a reciprocal impact on motility and biofilm formation (Xu *et al*., 2013). *visN* and *visR* mutants are non-motile but also over-produce the adhesive unipolar polysaccharide (UPP), which results in a hyper-attachment phenotype. This is despite the observation that non-motile and aflagellate *A. tumefaciens* mutants have a significant deficiency in surface attachment under non-flowing conditions, a phenotype likely due to inefficient engagement with surfaces (Merritt *et al*., 2007). The increased UPP production in *visN* and *visR* mutants is due to elevated levels of the intracellular second messenger cyclic diguanylate monophosphate (cdGMP) and is mediated through control of specific cyclic-di-GMP synthases (diguanylate cyclases, DGCs) (Xu *et al*., 2013). It is unclear to what extent the ExoR regulon intersects with the VisNR regulon to coordinate the complex developmental phenotypes of motility and biofilm formation.

We previously described transcriptome analyses for both the VisNR regulon (Xu *et al*., 2013) and the ExoR regulon (Heckel *et al*., 2014). The VisNR regulon was determined by comparing the transcriptomes of a Δ*visR* strain and wild-type *A. tumefaciens*, and the ExoR regulon was determined by comparing Δ*exoR* to wild-type. The VisNR and ExoR regulons shared a striking congruence of down-regulated genes in the motility and chemotaxis category; 51 genes from over a dozen operons display severely reduced expression in both analyses. This is not surprising for the VisNR regulatory network; VisNR are class IA motility regulator proteins, so we expect expression of all flagellar synthesis and chemotaxis genes to be dependent on VisN and VisR. Importantly, expression of *rem* (ATU_RS02820), encoding the class IB regulator was greatly reduced in the Δ*visR* mutant compared to wild-type. Although almost all the motility and chemotaxis genes are also dramatically decreased in the Δ*exoR* mutant, conspicuously absent are the class I motility regulators *visN, visR*, and *rem* (Heckel *et al*., 2014). These findings suggest that the ExoR-ChvG-ChvI pathway impacts motility downstream of *rem* expression. In this study, we examined the interaction between these pathways and discovered a previously undefined ExoR-ChvG-ChvI-regulated target gene that directly impacts Rem-dependent gene activation.

## Results

### Convergence of the VisN-VisR-Rem and ExoR-ChvG-ChvI Pathways

We previously demonstrated that deletion of the *chvI* gene (ATU_RS00165) relieves the inhibition of motility and chemotaxis genes in the Δ*exoR* (ATU_RS08400) mutant (Heckel *et al*., 2014). To further investigate the role of *chvI,* we examined the impact of mutating its site of phosphorylation. As with many two-component system response regulators, ChvI is predicted to be phosphorylated at a conserved aspartate residue (D52) in its N-terminal receiver domain to become active, in this case via the activity of the ChvG sensor kinase (Fig. S1) (Cheng & Walker, 1998). We mutated the *chvI* codon for this aspartate to glutamate (D52E) and to asparagine (D52N), to mimic and prevent phosphorylation by ChvG, respectively. Allelic replacement mutants of the native *chvI* gene with these alleles in the wild type and the Δ*exoR* mutant were then tested for control of motility and biofilm formation. Similar to Δ*exoR*, the *chvID52E* mutant was non-motile and had a severe biofilm defect; the phenotypes of this mutant were not impacted by deletion of *exoR* (Fig. 1A). In contrast, the *chvID52N* mutation had no effect on motility in an otherwise wild-type background but was completely epistatic to the non-motile and non-biofilm Δ*exoR* mutant phenotypes. The *chvID52E* mutant also phenocopies the Δ*exoR* deletion strain for elevated succinoglycan production (Heckel *et al*., 2014, Tomlinson *et al*., 2010), evidenced by dramatic binding of Calcofluor, a polysaccharide-specific dye, and diffuse fluorescence around the colony (Fig. S2). In contrast, when the native copy of *chvI* was replaced with the *chvID52N* allele, colonies show a slight decrease in Calcofluor binding relative to wild type (Fig S2). The Δ*exoR* Δ*exoA* (ATU_RS18935) double mutant does not produce succinoglycan, and does not stain with Calcofluor (Tomlinson, 2010 #1702).

**Figure 1.**
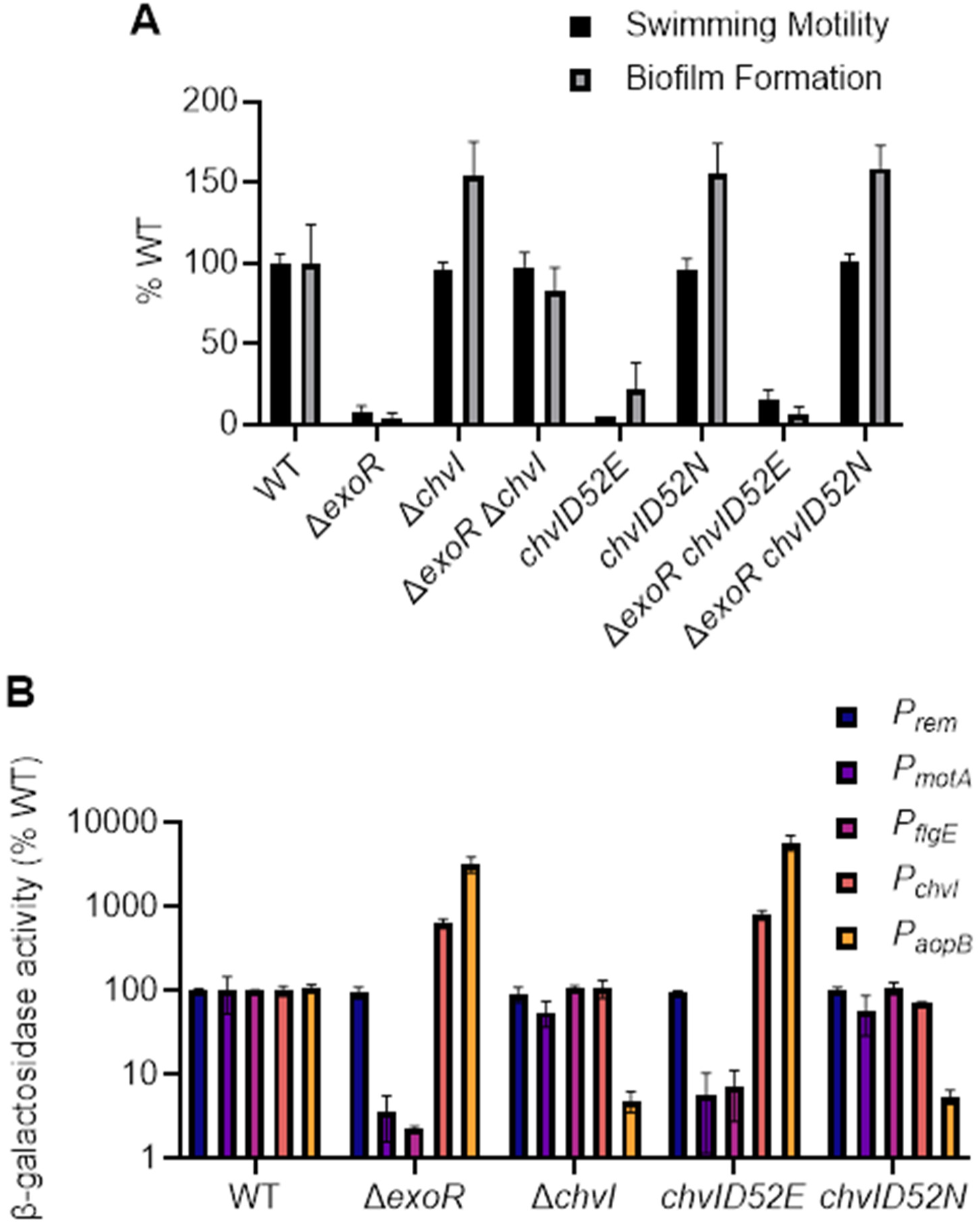
Phenotypes from *chvID52E* and *chvID52N* mutations. (A) The swimming and biofilm deficiencies of the Δ*exoR* mutant are recapitulated by the ChvID52E allele. Motility was assessed as swim ring diameter in 0.3% Bacto Agar after three days of incubation in a humid chamber at room temperature (RT) and reported values are relative to wild-type C58. Biofilms were developed on PVC coverslips in ATGN media supplemented with 22 μM FeSO_4_・7H_2_O for 48 h. Biofilm biomass was determined by solubilizing 0.1% crystal violet-stained biofilm cells in 33% acetic acid and measuring the absorbance at 600 nM (A_600_). Crystal violet absorbance was normalized per cell count by dividing by the planktonic culture OD_600_. The data are presented as biofilm biomass relative to wild type. Error bars represent standard deviation for at least three replicates for each strain. (B) β-galactosidase assays of *A. tumefaciens* derivatives harboring plasmid-borne *lacZ* fusions. Activity of the five indicated promoters was assessed by translational fusions to *lacZ* and measurement of β-galactosidase activity.. Cultures were inoculated from colonies and grown to exponential phase (OD_600_ 0.3 – 0.8) for promoter activity assessment. Error bars represent standard deviation for at least three replicates.

Deletion of *exoR* does not alter expression of a plasmid-borne *rem*-*lacZ* translational fusion (Fig. 1B) nor does it impact a similar *lacZ* translational fusion to *visN* (Fig. S3). These results are consistent with transcriptomic analysis of the Δ*exoR* mutant, as there is no substantial reduction of *visN, visR,* or *rem* transcript abundance in this genetic background (Heckel *et al*., 2014). In fact, *rem-lacZ* expression remains unchanged in the Δ*chvI*, *chvID52E*, and *chvID52N* mutants (Fig. 1B). In contrast, similarly constructed *lacZ* fusions to the motility genes *motA* (ATU_RS02760) and *flgE* (ATU_RS02825), encoding a motor protein and the flagellar hook protein respectively, were greatly diminished in the Δ*exoR* and *chvID52E* mutants, but equivalent to wild type in the Δ*chvI* null and *chvID52N* mutants. Conversely, activity of *lacZ* fusions to *chvI* itself and the predicted outer membrane protein *aopB* (ATU_RS05585) were strongly elevated in Δ*exoR* and *chvID52E* mutants (Fig. 1B); these results correlate well with prior transcriptomic data suggesting that the transcription of *chvI* and *aopB* are elevated in the Δ*exoR* mutant (Heckel *et al*., 2014). The *aopB* fusion was decreased relative to wild type in Δ*chvI* and *chvID52N* mutants, whereas the *chvI-lacZ* was equivalent to wild type. Thus, although these data suggest that phospho-ChvI activates expression of target genes such as *aopB*, it inhibits motility gene expression, and does so independent of changes to *rem or visNR* expression. Therefore, we explored other models of how ChvI might be regulating motility gene expression.

### ChvI and Rem independently bind their own target promoters but do not cross-recognize binding sites and do not interact with each other *in vitro*

An alternate model for ChvI-mediated inhibition of flagellar motility gene expression was that ChvI directly competes with Rem for binding to Rem-dependent motility promoters thereby preventing their transcriptional activation. ChvI is a PhoB/OmpR class response regulator, with a winged helix-turn-helix motif, and it regulates many genes in *A. tumefaciens* (Heckel *et al*., 2014, Okamura *et al*., 2000). Prior studies demonstrated binding of purified ChvI^D52E^ protein *in vitro* to the promoter region of the type VI secretion system (T6SS) genes, specifically the intergenic region between the divergent operons that initiate with *impA* (ATU_RS20340) and *clpV* (ATU_RS20345) (Wu *et al*., 2012). We tested the ability of purified preparations of N-terminal, hexahistidinyl-tagged derivatives of ChvI^D52E^ and wild type ChvI to bind to a PCR amplicon of this promoter region (defined as *P_hcp_*) *in vitro* as determined by electrophoretic mobility shift assay (EMSA) (Fig. 2A-B). Consistent with the prior study, His_6_-ChvI^D52E^ was able to robustly shift the *P_hcp_* DNA fragment (Fig. 2A). In contrast to the prior study, however, we observed similar binding of *P_hcp_* by the wild-type His_6_-ChvI allele, presumably the non-phosphorylated form of the protein (Fig. 2B). However, addition of either protein does not shift the mobility of DNA fragments from any of the upstream regions of motility genes (data not shown). To determine more specifically where ChvI binds the *hcp* promoter, we created smaller fragments of different portions of *P_hcp_* (Fig. S4). We found that at least 140 bp upstream of the start codon of *clpV* were required in the amplicon to observe a gel mobility shift following incubation with His_6_-ChvI^D52E^, well beyond the transcription start site we mapped by 5’ RACE to be 63 bp upstream of the start codon of *clpV.* We also used 5’ RACE to map the transcription start sites of *chvI* and *aopB*, and found them to start 84 and 142 bp upstream of the start codon, respectively. However, we did not observe His_6_-ChvI^D52E^ binding to these promoters by EMSA (data not shown).

**Figure 2.**
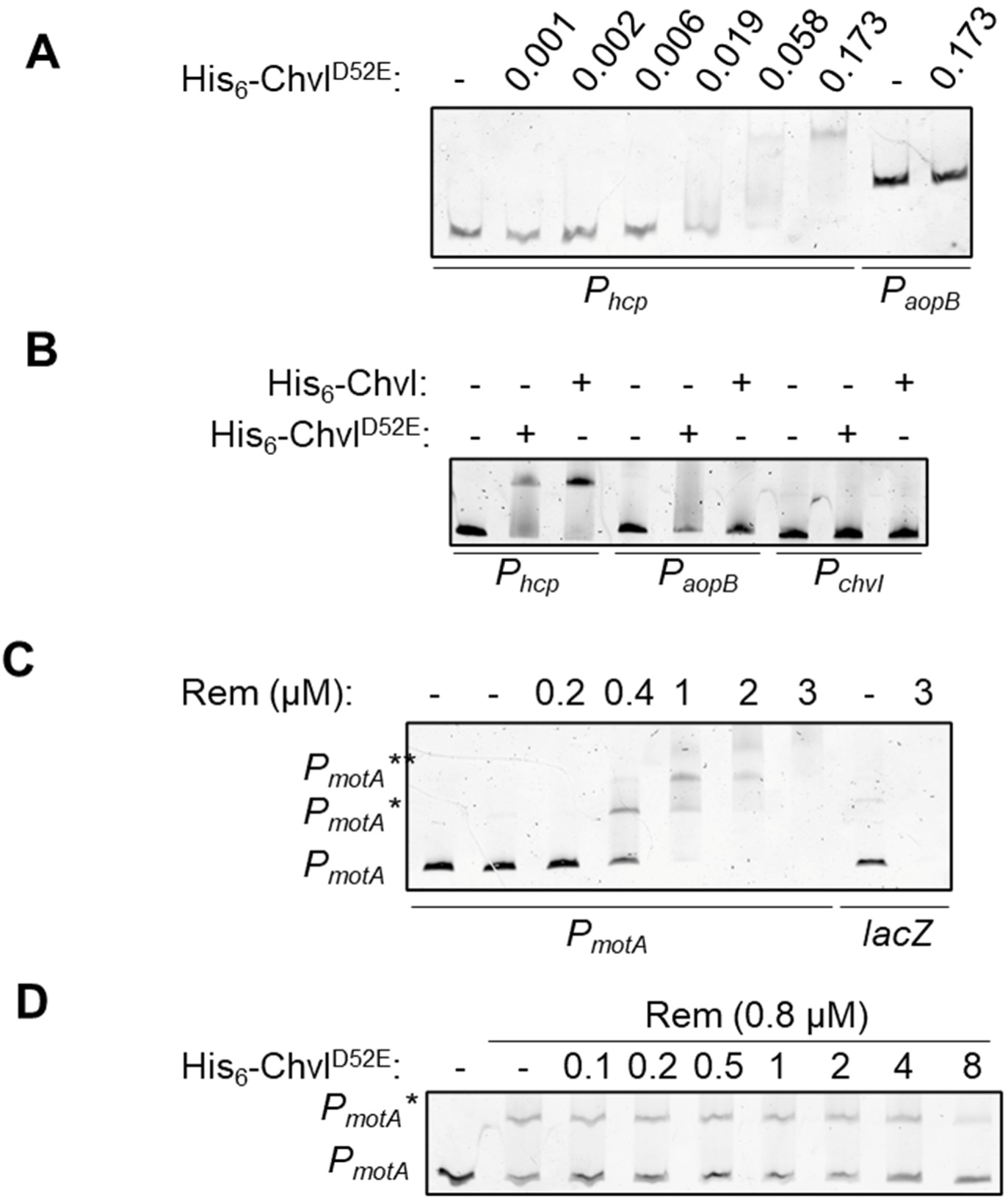
Electrophoretic mobility shift assay (EMSA) of DNA binding by ChvI and Rem. (A) Electrophoretic mobility shift assays were performed to evaluate binding of His_6_-ChvI^D52E^ to the intergenic region between the divergent *P_tss_* and *P_hcp_* operons. DNA containing putative binding sites (10 nM) was incubated with increasing concentrations of purified His_6_-ChvI^D52E^. Binding was assessed by DNA migration of reactions run on a 6% polyacrylamide native gel. DNA was detected by staining with SYBR Safe dye. One discrete shift (indicated by asterisk) is observed when DNA is incubated with purified His_6_-ChvI^D52E^. The *aopB* promoter was used as a negative control for binding. (B) DNA binding of His_6_-ChvI^D52E^ and His-ChvI (wild-type) alleles. Upstream DNA of *hcp, aopB, and chvI* (10 nM) was incubated with and without 600 nM of purified His-tagged alleles of ChvI and assessed for binding. (C) Binding of Rem to 10 nM fragment with *P_motA_* (D) His_6_-ChvI^D52E^ does not compete for Rem-*P_motA_* association. His_6_-ChvI^D52E^ was incubated with Rem for 10 min prior to the addition of 10 nM DNA to reactions and 20 min incubation prior to running samples on a 6% acrylamide gel and staining with SYBR Safe dye.

Rem is a two-component-type response regulator, with no known cognate sensor kinase, and it initially seemed plausible that ChvG might act to phosphorylate Rem as well as ChvI (although that model would not readily explain the impact of the *chvID52E* mutant on motility gene expression). Alignment of Rem with well-characterized response regulators indicates that, at the position for the conserved aspartic acid that is the phosphorylation site of canonical response regulators, Rem instead has a glutamate residue at this position (E50; Fig. S1). However, there is an aspartate five residues more N-terminal to this site (D45). To test whether these residues are important for Rem function, we expressed *rem* with the glutamate (RemE50N) or with the upstream aspartate residue (RemD45N), mutated to an asparagine. Expression of the RemE50N allele complemented a Δ*rem* mutant for motility (Fig S5A) and biofilm formation (Fig S5B) similarly to the wild-type *rem* allele and had no impact on the non-motile *exoR* phenotype. The RemD45N allele partially restored motility and biofilm formation to the Δ*rem* mutant. Together, these results suggest that the native residues D45 and E50 are not required for the ability of Rem to bind the class II motility promoters, although the D45N mutation may partially compromise Rem activity.

The Rem protein belongs to the emerging family of aspartate-less receiver domain (ALR)-containing response regulators, which have receiver domains lacking the canonical aspartate residue that is typically the site of phosphorylation leading to modulation of the protein’s activity (Maule *et al*., 2015). A subset of this protein family contain a single cysteine residue in this or nearby position. One such ALR-containing protein, RitR from *Streptococcus pneumoniae,* contains a single cysteine residue in the linker region between the receiver and DNA-binding domain and requires the cysteine residue for redox state-dependent dimerization, thereby impacting gene regulation (Glanville *et al*., 2018). Mutation of the cysteine to serine or aspartate simulates a reduced or oxidized state, respectively and modulates DNA-binding activity. Rem contains a single cysteine at position 143 and we hypothesized that Rem might function similarly (Fig. S1). However, expression of *rem* alleles with Rem143C mutated to serine, aspartate or alanine complemented motility equivalently to the wild-type allele (Fig. S6). The cysteine residue in this position is also not conserved in Rem orthologs (Fig. S1).

The transcriptional start sites of Rem-regulated class II flagellar genes *_flgB_* (ATU_RS02735) and *_motA_* (ATU_RS02760) were determined by 5’ RACE mapping (Rapid Amplification of cDNA ends) and found to be downstream of an imperfect direct repeat sequence that is conserved among multiple *A. tumefaciens* predicted class II motility gene promoters (**CG**-W**CAAG**WCTCR**CG**-**CAAG**NYYNN**AC**; Fig. S7A) and similar to repeats bound by Rem upstream of motility genes in *S. meliloti* (Rotter, 2006 #1465). The divergent motility gene operons that initiate with ATU_RS02755 and *motA* have two of these direct repeat sequences in their intergenic region. Rem was purified using an Intein tag system (in which the tag self processes to release a nearly native protein). When complexed with PCR-amplified *P_motA_,* Rem forms two shifted EMSA complexes (Fig. 2C), indicative of two Rem binding sites. The *P_flgB_* fragment only contains a single putative Rem binding site and forms a single Rem-DNA complex (Fig. S7B), which suggests that Rem binds one site at this promoter. We did not observe binding of Rem to *P_rem_* (Fig. S7C). To test whether ChvI competes with Rem for DNA binding, we incubated purified His_6_-ChvI^D52E^ with Rem and the *P_motA_* amplicon. We assessed binding by mixing all three components (Rem, His_6_-ChvI^D52E^, and *P_motA_)* simultaneously and also with addition of His_6_-ChvI^D52E^ to pre-formed Rem-DNA complexes. We did not observe specific binding of His_6_-ChvI^D52E^ to *P_motA_,* nor did His_6_-ChvI^D52E^ disrupt Rem-*P_motA_* complexes (Fig. 2D, S8A) other than at very high protein concentrations that led to non-specific, large aggregates.

Given that ChvI and Rem are both response regulators, we hypothesized that ChvI might directly interact with Rem. To test for interaction between these proteins, we performed farwestern assays, immobilizing Rem on nitrocellulose membranes and probing with His_6_-ChvI^D52E^. Upon development with monoclonal anti-His_6_ antibody, we did not observe stable association of ChvI and Rem (Fig. S8B). Overall, the genetic interaction we have observed between the ExoR-ChvG-ChvI and the VisNR-Rem pathways could not be explained by physical interaction between ChvI and Rem or ChvI and the target promoters of Rem. We therefore hypothesized that a previously-unidentified ChvI-regulated factor inhibits Rem-dependent motility gene activation upon growth in acidic medium or in upon deletion of *exoR*.

### Isolation of motile suppressors of Δexo*R*

To identify a cellular component that mediates the motility inhibition by ChvI, we performed a transposon-mediated motility suppressor screen to identify mutants that restore motility to a Δ*exoR* Δ*exoA* double mutant (*exoA* is mutated to prevent the mucoidy associated with succinoglycan overproduction in the Δ*exoR* mutant) (Heckel *et al*., 2014). For simplicity, we will list the Δ*exoR* Δ*exoA* double mutant strain as Δ*exoR* unless otherwise specified. As observed before and therefore expected, multiple transposon insertional mutations were identified in the *chvI*-*chvG* locus. In addition to reversing motility, these null mutations also rescue the biofilm deficiency of Δ*exoR* (Heckel *et al*., 2014). Expression of the *chvID52N* allele fails to correct the biofilm formation of Δ*exoR* (Fig. 1A), indicating that the biofilm deficiency of Δ*exoR* is dependent on the phosphorylation of ChvI. Although flagellar motility is correlated with efficient biofilm formation in *A. tumefaciens* (Merritt *et al*., 2007), the biofilm deficiency of the Δ*exoR* mutant is more severe than that for the non-motile mutants such as Δ*rem,* indicating that there are additional functions besides motility under ExoR control that impact biofilm formation (Fig. S9). We screened for Δ*exoR* suppressor mutants that rescued the motility defect but retained the biofilm deficiency of Δ*exoR*, and therefore retained native *chvG-chvI.* We obtained three such motile, biofilm-deficient transposon mutants BCH132, BCH133, and BCH134 (Fig. 3A, Fig. S10A). To confirm that the suppressor mutations do not alter the activity of ChvG-ChvI, we assessed expression of the ChvI-regulated genes *chvI* and *aopB* through activity of *lacZ* translational fusions in BCH133 (Fig. S10B). We found that BCH133 retained the elevated *P_chvI_* and *P_aopB_* activity indicative of active of ChvG-ChvI. Since Δ*exoR* has a nonmotile phenotype but does not alter *rem* expression (Heckel *et al*., 2014), we hypothesized that these suppressors of Δ*exoR* would restore motility without changing *rem* expression. Indeed, we observed no change in *P_rem_* activity in BCH133 relative to its Δ*exoR*Δ*exoA* parent strain (Fig. S10B). Sequencing at the transposon junctions revealed BCH132 had a transposon insertion within ATU_RS03690, a *bioY* homolog likely involved in biotin biosynthesis. The transposon in BCH133 mapped to ATU_RS10935, which is predicted to code for an ABC transporter subunit. BCH134 carried an insertion within ATU_RS09055, which is homologous to *bolA* from *E. coli,* which negatively regulates motility and positively regulates biofilm formation (Dressaire *et al*., 2015). Although we hypothesized that BolA may regulate these phenotypes in *A. tumefaciens,* deletion of *bolA* in the Δ*exoR* mutant did not restore motility or biofilm formation, nor did the deletion independently affect these phenotypes (Fig. S11A-B). Since all three transposon mutants had similar phenotypes with unrelated transposon insertion sites, we subjected BCH132-4 to whole-genome resequencing. The single mutations from all three isolates clustered to the same general genomic location, diagrammed in Fig 3A and were first mapped to the initial *A. tumefaciens* C58 reference genome. The independent mutations included a single base substitution (BCH134), a base deletion in the 3’ end of the annotated hypothetical gene Atu1638 (BCH133), and a single base substitution in the intergenic region between Atu1638 and the adjacent gene Atu8018 (ATU_RS08045) (BCH132). We recreated the *mirA* frameshift suppressor mutation of BCH133 in a naïve Δ*exoR* background (Δ*exoR mirA*_FS_); the mutation restored motility to Δ*exoR* (Fig. S10C) but retained the biofilm defect of the parent strain (data not shown). The point mutation had no effect on swimming motility in the wild-type background, suggesting that motility may be saturated at wild type levels, or that the effects of this mutation are specific to an Δ*exoR* background. Given the proximity of the mutations and the fact that they all shared the same phenotype, the locus was investigated further. Introduction of an in-frame stop codon after amino acid 19 (serine) within the Atu1638 coding sequence did not restore motility to *ΔexoR* (Fig. S12A), suggesting that Atu1638 may not be required for the motility defect of *ΔexoR*. A survey for over-annotated genes performed by Yu et al. predicted that Atu1638 was erroneously annotated as a protein-coding gene (Yu *et al*., 2015), and Atu1638 is not annotated in a more recent C58 genome annotation.

**Figure 3.**
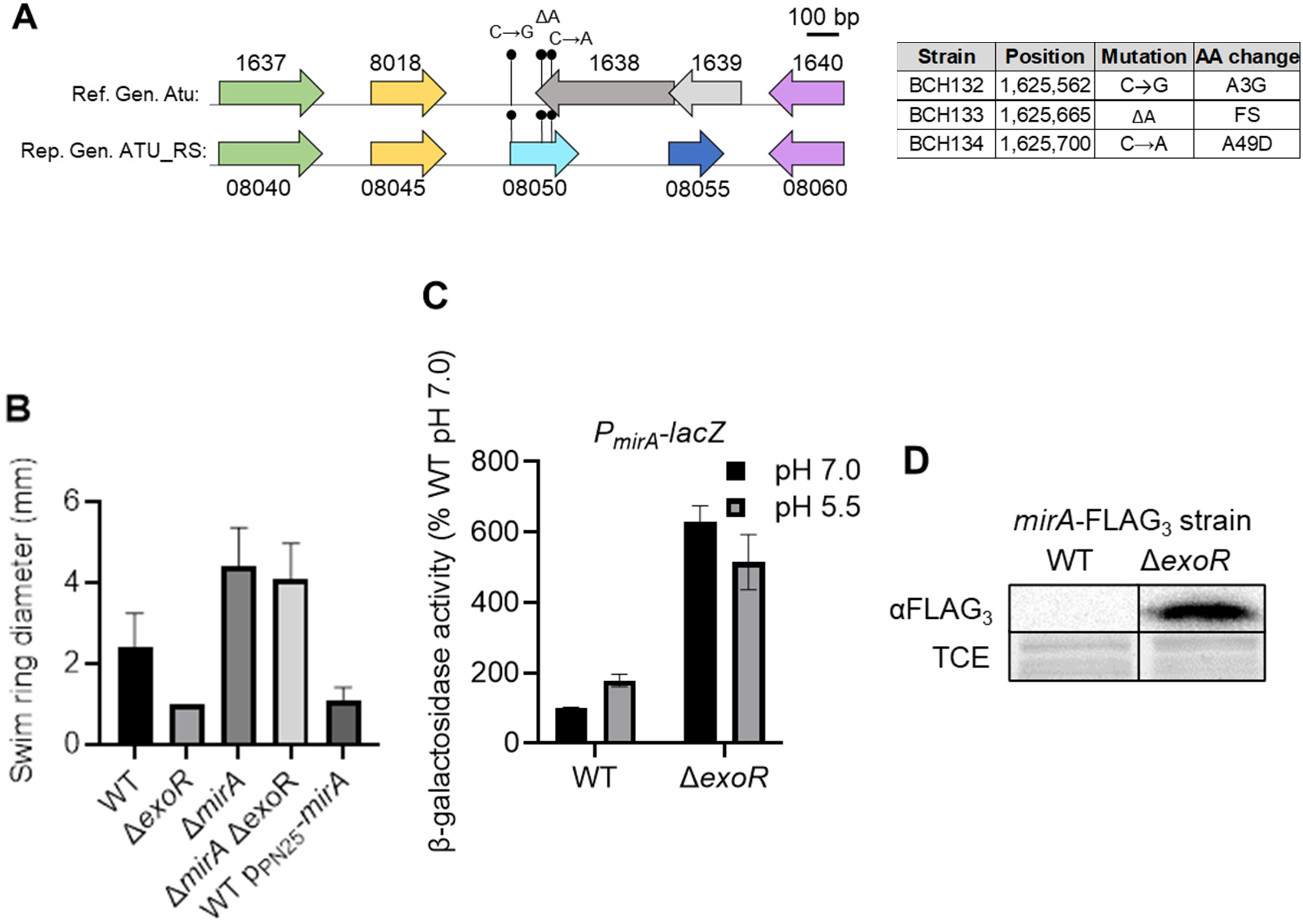
Identification of the motility inhibitor gene *mirA*. (A) Top arrows: gene diagram with Atu gene numbers (labeled above gene arrow) from the C58 reference genome (GenBank access. no. AE008687 for the circular chromosome) (Goodner *et al*., 2001). Bottom arrows: gene diagram with ATU_RS numbers (labeled above arrows) from the C58 representative genome annotation from 03-10-2020 (Genbank access. no. (NCBI-RefSeq:NC003062 for the circular chromosome). The direction of arrows indicates the direction of transcription. Stemmed circles indicate mutation sites with mutations are labeled above. The table on the right notes the genomic location (nucleotide number on C58 circular chromosome) and the resulting amino acid change for ATU_RS0850 is listed as AA change for each mutation. (B) Motility assay of *mirA* mutation in wild-type and Δ*exoR (ΔexoA*) strains as well as wild type expressing p *P*_N25_-*mirA.* Colonies were inoculated into the center of 0.3% Bacto Agar and incubated for 24 h in a humid chamber at RT prior to swim ring measurement. Error bars represent standard deviation for at least three replicates. (C) Activity of the *P_mirA_*-*lacZ* fusion following growth in neutral conditions (ATGN pH 7) or acidic medium (AT-MES pH 5.5) in wild-type and Δ*exoR (ΔexoA*) strains. Strains were grown to exponential phase prior to freezing at −20 °C and thawing prior to β-galactosidase assays. Error bars represent standard deviation for at least three replicates. (D) Western blot analysis of *mirA*-FLAG_3_ protein levels. Strains with the native *mirA* locus replaced with *mirA*-FLAG_3_ were grown to exponential phase prior to separation of whole cell lysates by SDS-PAGE (12.5% polyacrylamide gel) and transfer to a nitrocellulose membrane and probing with anti-M2 antibody. Images are from a single western blot of a single gel, with identical image treatment. Bands from total protein stained with 2,2,2 trichloroethanol (TCE) are shown as a loading control.

Closer observation of this locus revealed that a small open reading frame (231 bp) overlaps all three mutations; this reading frame was not annotated in the *A. tumefaciens* C58 reference genome, but is annotated by the MicroScope database as AERS4k1_1608 (Vallenet *et al*., 2009), and in the recent re-annotation of the *A. tumefaciens* C58 genome as ATU_RS08050. We hypothesized that the protein product of ATU_RS08050 may be required for the gain of motility in our isolated suppressors of Δ*exoR.* As detailed below, we have now designated this gene as the *mirA.* To test whether *mirA* is necessary for the non-motile phenotype of Δ*exoR* mutants, we constructed an in-frame deletion mutant in both the wild type and Δ*exoR* backgrounds. We observed wild-type motility in both the Δ*mirA* and Δ*exoR* Δ*mirA* strains (Fig. 3B), suggesting that this gene is necessary for motility inhibition by the ExoR-ChvG-ChvI system. Given our prior observations that the ExoR-ChvG-ChvI pathway inhibits expression of class II flagellar genes (Heckel *et al*., 2014), that are activated by Rem (Rotter *et al*., 2006), we tentatively designated the gene as *mirA*, for <u>m</u>otility inhibitor of <u>R</u>em. To test whether translation of *mirA* is required for the non-motile phenotyoe of *ΔexoR*, we replaced the start codon ATG with a nonfunctional ATC; this mutation suppressed the *ΔexoR* motility defect (Fig. S10C). Additionally, introduction of the amber stop codon TAG into the *mir*A reading frame (following glutamic acid 20, outside of the overlap with the initially annotated Atu1638 coding sequence) also suppressed the *ΔexoR* motility defect (Fig. S12B), indicating that translation of the *mirA* gene product is specifically required for motility inhibition in *ΔexoR* mutants.

### Expression of *mirA* is sufficient to inhibit motility and is under ExoR-ChvG-ChvI control

Multiple deep sequencing studies have identified a small RNA adjacent to, and overlapping with the *mirA* gene (1,625,555-1,625,785 bp on the C58 circular chromosome); these sRNAs detected by separate studies are *RNA353* (140 bp, from position1,625,422-1,625,562), an unnamed sRNA of 214 bp (from position 1,625,425-1,625,639, and C1_1625426F (469 bp, from position 1,625,426-1,625,895), the last of which fully overlaps the *mirA* gene (Dequivre *et al*., 2015, Lee *et al*., 2013, Wilms *et al*., 2012). In the study by Lee, et al., C1_1625426F was predicted to encode a small open reading frame, which, in this study, we are naming *mirA.* To test whether expression of *mirA* is sufficient to achieve motility inhibition, we ectopically expressed a plasmid-borne copy of the *mirA* coding sequence from the constitutive coliphage T5 promoter *P*_N25_ (Wang *et al*., 2000). We found that ectopic expression of the *mirA* coding sequence strongly inhibited motility to an equal extent as observed for the Δ*exoR* mutation (Fig. 3B), indicating that *mirA* is sufficient for motility inhibition. Plasmid-borne expression of *mirA* from its native promoter (including the *RNA353* sequence) (p*P_mirA_*::*mirA*) also showed a similar strong motility inhibition in a wild-type background (Fig. S13),. Expression from this plasmid also complemented the Δ*exoR* Δ*mirA* strain, bringing motility back to the level of the Δ*exoR* strain.

To determine whether the expression *mirA* is dependent on the ExoR-ChvG-ChvI pathway, we fused the promoter and start codon of *mirA* to *lacZ* and performed a β-galactosidase assay at pH 7 and 5.5. We found that activity of *P_mirA_*-*lacZ* was elevated in the Δ*exoR* mutant background and to a lesser extent in cells grown in pH 5.5 medium relative to pH 7 medium (Fig. 2C). Although we did not observe a change in *mirA* expression in our previous transcriptomic analysis of Δ*exoR* compared with wild type (Heckel *et al*., 2014), this is unsurprising as *mirA* was not previously annotated as a gene and was not included in the gene array that was used in that study. Expression of *mirA* was not affected by a frameshift mutation within the native *mirA* gene (Fig. S10D), indicating that *mirA* is not autoregulatory. To test for regulation of MirA protein levels, we replaced the native copy of *mirA* with a C-terminal, in-frame FLAG_3_ tag fusion. We found that levels of MirA-FLAG_3_ were elevated in Δ*exoR* (Fig. 2D), further evidence that MirA is regulated by the ExoR-ChvG-ChvI system.

### MirA regulates motility gene expression

Combining our data that *mirA* was necessary for motility inhibition in Δ*exoR* and that *mirA* gene expression is regulated by the ExoR-ChvG-ChvI system, we hypothesized that cells expressing *mirA* would be decreased for class II motility gene expression relative to wild type. To test our hypothesis that MirA regulates class II motility gene expression and to uncover genes regulated by MirA, we performed RNA-Seq on total RNA extracted from wild type cells harboring either an empty vector or the same plasmid with *mirA* expressed from the *P_mirA_* native promoter. A total of 131 genes were differentially expressed (log_2_ fold change <-0.50 or >0.50 and p<0.05)(Table S1, Fig. 4). The majority of these genes had decreased expression in cells ectopically expressing *mirA* (112 genes), and we observed elevated expression of a much smaller subset of the genes (19 genes). In strains harboring the *mirA* plasmid, expression of genes of the flagellar gene clusters and chemotactic genes were strongly down-regulated (Table S1; Fig. 4, blue circles). Since Δ*exoR* mutants are non-motile but retain wild-type expression of flagellar gene activators *visNR* and *rem* (Fig. S3) (Heckel *et al*., 2014), we hypothesized that strains with elevated *mirA* expression would not lead to changes in expression of these genes. Indeed, we observed that expression of *visN*, *visR*, and *rem* were not significantly altered by expression of *mirA* (Table S1). Comparison of the transcriptomes from Δ*exoR* vs wild type with the derivative with ectopic *mirA* expression reveals that most genes in the MirA regulon are a subset of genes regulated by ExoR and VisR, with 96 genes differentially regulated when comparing Δ*exoR* and/or Δ*exoR* to wild type (log_2_ fold change <-0.50 or >0.50 and p<0.05)(Table S2)(Heckel *et al*., 2014, Xu *et al*., 2013). A gene cluster that was uniquely down-regulated in the *mirA* dataset was a predicted prophage, spanning from Atu1183 – Atu1194 (ATU_RS05840-ATU_RS05900). However, expression of these genes was extremely low in cells harboring both p*mirA* and the empty vector, so the importance and relevance of this cluster is unknown. Expression of the T6SS genes *icmF* and *impI* (ATU_RS020285 and ATU_RS020300, respectively) was slightly elevated in the strain harboring the *mirA* plasmid, and expression of these genes is also elevated in the Δ*exoR* mutant (Heckel *et al*., 2014). As we have demonstrated that His_6_-ChvI^D52E^ binds at the promoter adjacent to this locus (*P_hcp_)*(Fig. 2B), it is surprising that MirA may also regulate expression of these genes. Mutation of *mirA* does not affect *chvI* expression (Fig. S10B), so the increase in T6SS gene expression is unlikely to be due to elevated *chvI* expression upon *mirA* induction. Similarly, the increase in expression of ATU_RS20300 cannot simply be explained as a decrease in Rem-dependent gene activation because expression of ATU_RS20300 is decreased in a Δ*visR* mutant, where *rem* is not expressed (Heckel *et al*., 2014).

**Figure 4.**
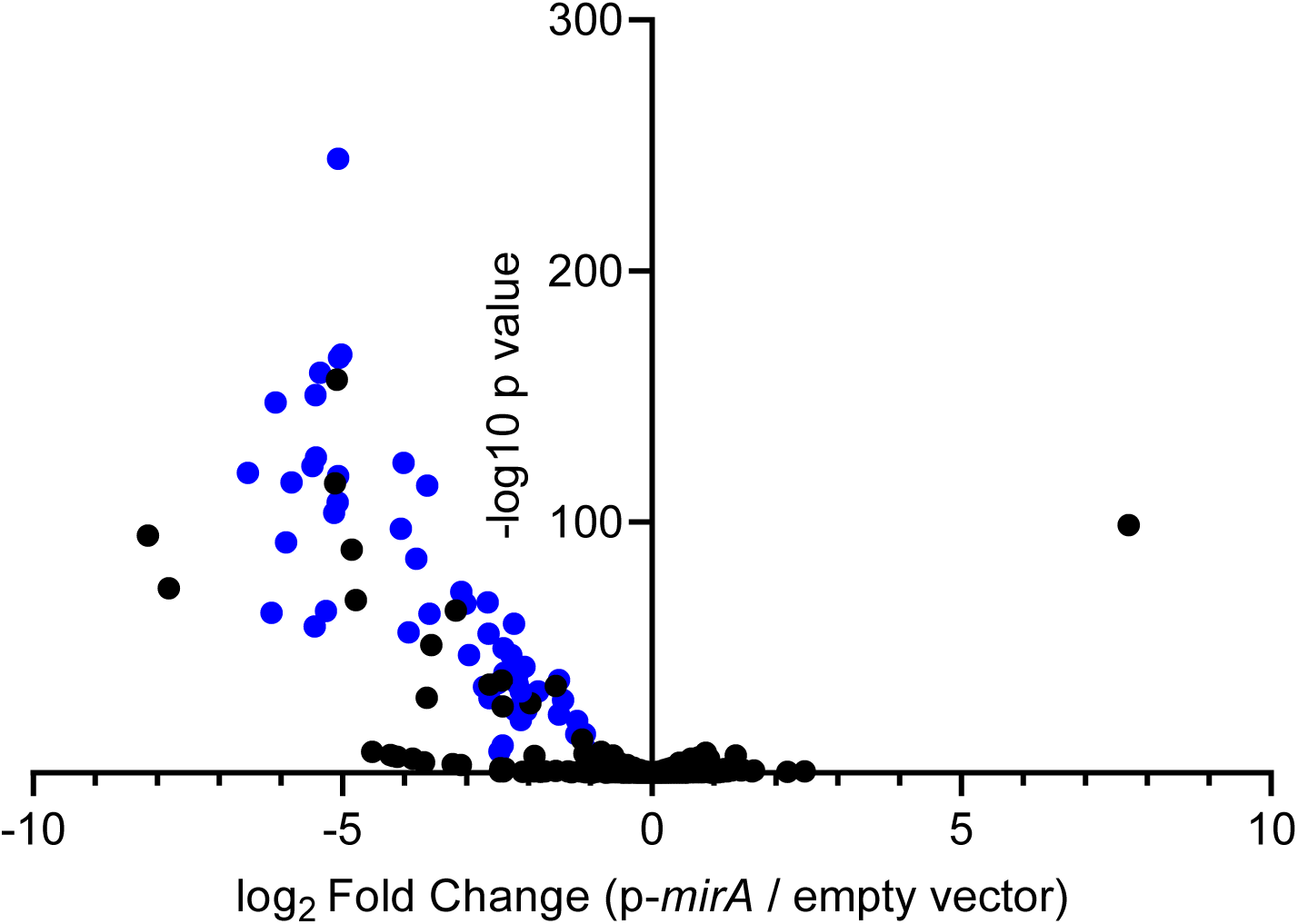
Ectopic *mirA* expression leads to the downregulation of motility and chemotaxis genes. Log_2_(fold change) and −log_10_(p value) values for all *A. tumefaciens* genes from RNAseq of wild type C58 harboring an empty vector (pSRKGm) or p*mirA* (pMAT14, p*P_lac_*::*P_mirA_*::*mirA*), with three replicates per plasmid. Blue shaded dots label genes belonging to flagellar or chemotaxis gene clusters.

### MirA interacts with Rem *in vitro*

We hypothesized that MirA dampens motility gene expression through sequestration of the motility activator protein Rem. To test whether MirA interacts directly with Rem, we performed a pull-down assay with purified untagged Rem and MirA fused to a C-terminal hexahistidinyl tag (MirA-His_6_). We used cobalt resin, which interacts strongly with hexahistidinyl tags (Lichty *et al*., 2005), for a pull down assay. Rem did not bind the cobalt resin alone, but co-eluted with MirA-His_6_ following co-incubation of the proteins (Fig. 5A). We also performed a farwestern experiment to separately test MirA-Rem interaction. Dilutions of purified Rem were separated by SDS-PAGE and transferred to a nitrocellulose membranes, and following blocking, these were incubated with blocking agent either with or without purified MirA-His_6_. Bound MirA-His_6_ was detected with anti-His_6_ monoclonal antibodies. The strength of anti-His_6_ chemiluminescent signal correlated with the concentration of Rem protein and depended on the presence of MirA-His_6_ (Fig. 5B).

**Figure 5.**
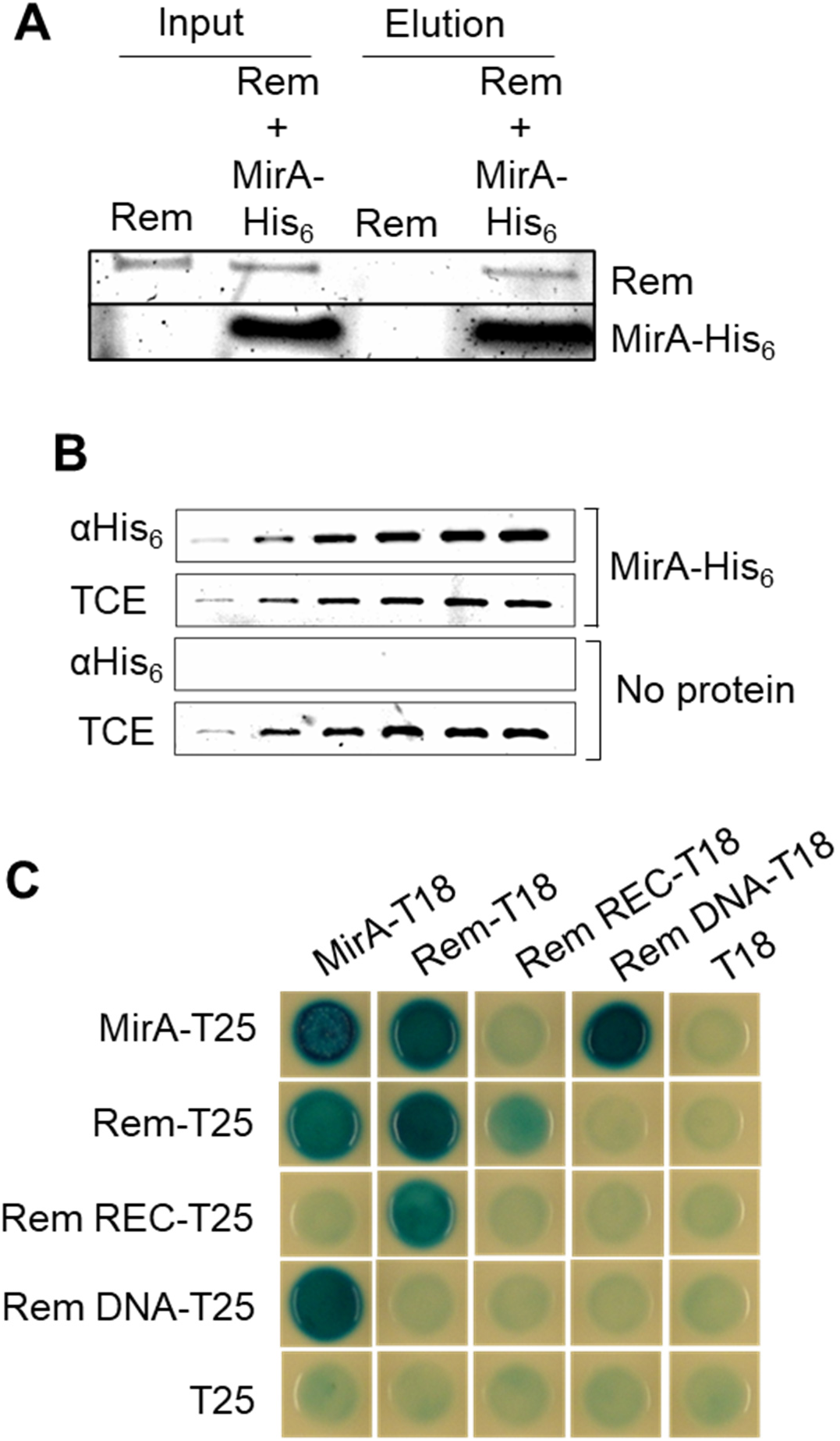
MirA directly interacts with the Rem DNA-binding domain. (A) Pull down assay reveals interaction between Rem and MirA-His_6_. Rem was incubated alone or with MirA-His_6_ for 30 min prior to addition of the samples to 10 μL of TALON cobalt resin. Following a 30 min incubation and three washes to remove protein that fails to bind the resin, samples were eluted in SDS-PAGE loading buffer and aliquots were electrophoresed on a Tris-Tricine gel labeled with 2,2,2 trichloroethanol (TCE). (B) Farwestern analysis of Rem and MirA-His_6_. Purified Rem protein was separated by SDS-PAGE (12.5% acrylamide) and immobilized on a nitrocellulose membrane and probed with MirA-His_6_ (or no protein as a control) and incubated with monoclonal anti-His_6_ antibody. Images of SDS-PAGE gels stained with 2,2,2 trichloroethanol (TCE) are shown as loading controls. (C) Bacterial Adenylate Cyclase Two-Hybrid (BACTH) assay of MirA and Rem. MirA and Rem as well as the Rem receiver and DNA-binding domains were fused to the T25 and T18 domains of *E. coli* adenylate cyclase for BACTH and assayed for *lacZ* activity when co-expressed in *E. coli* by plating 2 μL on LB agar with X-gal (160 μg/mL) and IPTG (500 μM) following a 24 h incubation.

### Bacterial two-hybrid analysis of MirA-Rem interactions

We used the Bacterial Adenylate Cyclase Two-Hybrid (BACTH) assay to further assess interaction between Rem and MirA fusion proteins expressed in *E. coli*. This assay detects protein-protein interactions that bring the T25 and T18 fragments of adenylate cyclase into close enough proximity to become catalytically active to synthesize cyclic AMP (cAMP) synthesis (Battesti & Bouveret, 2012). cAMP synthesis activates expression of *lacZ* which leads to blue colonies in the presence of X-gal. Co-expression of the MirA-T25 and Rem-T18 fusions resulted in blue colonies, further supporting a direct interaction between MirA and Rem (Fig. 5C). A similar result was obtained for the reverse combination Rem-T25 and MirA-T18, while negative controls harboring a sole MirA- or Rem-adenylate cyclase fragment yielded negative results, indicated by white colonies. Strains expressing MirA-T25 and MirA-T18 or Rem-T25 and Rem T18 also resulted in blue colonies, suggesting self-interactions of MirA and Rem, in addition to their formation of a MirA-Rem heterocomplex. As with many two-component-type response regulator transcription factors (Gao *et al*., 2007), the Rem protein contains a receiver domain and DNA-binding domain. To assess which domain of Rem interacts with MirA, we tested the receiver and DNA-binding domains individually for interaction with MirA via BACTH. We observed a strong interaction of MirA with the DNA-binding domain of Rem, and little to no interaction of MirA and the receiver domain (Fig. 5C).

### MirA inhibits Rem-DNA binding

Given our BACTH data supporting a model where MirA interacts with the Rem DNA-binding domain, we hypothesized that this interaction might impair Rem DNA-binding activity. To test this model, we co-incubated purified Rem and MirA-His_6_ protein prior to incubation with a target promoter of Rem, *P_motA_*, and performed electrophoretic mobility shift assays (Fig. 6A). Co-incubation with increasing concentrations of MirA-His_6_ led to disruption of Rem-DNA binding, even at molar ratios of the two proteins at or below one-to-one. To test whether MirA-His_6_ could disrupt pre-formed complexes between Rem and *P_motA_* promoter DNA, we incubated Rem with the promoter DNA to allow complexes to form, and then added MirA-His_6_ at varying concentrations. As the concentration of MirA-His_6_ increased, we observed an increase in free DNA relative to the Rem-DNA complex (Fig. 6B). To test whether DNA-binding inhibition by MirA was specific to Rem, we co-incubated MirA-His_6_ with His_6_-ChvI^D52E^ and assessed DNA binding by ChvI (Fig. S14). Even at concentrations of MirA-His_6_ that inhibited Rem-DNA binding activity, His_6_-ChvI^D52E^ was still able to bind the *P_hcp_* promoter DNA, suggesting that MirA inhibition of DNA binding is specific to Rem and not an inherent property of MirA.

**Figure 6.**
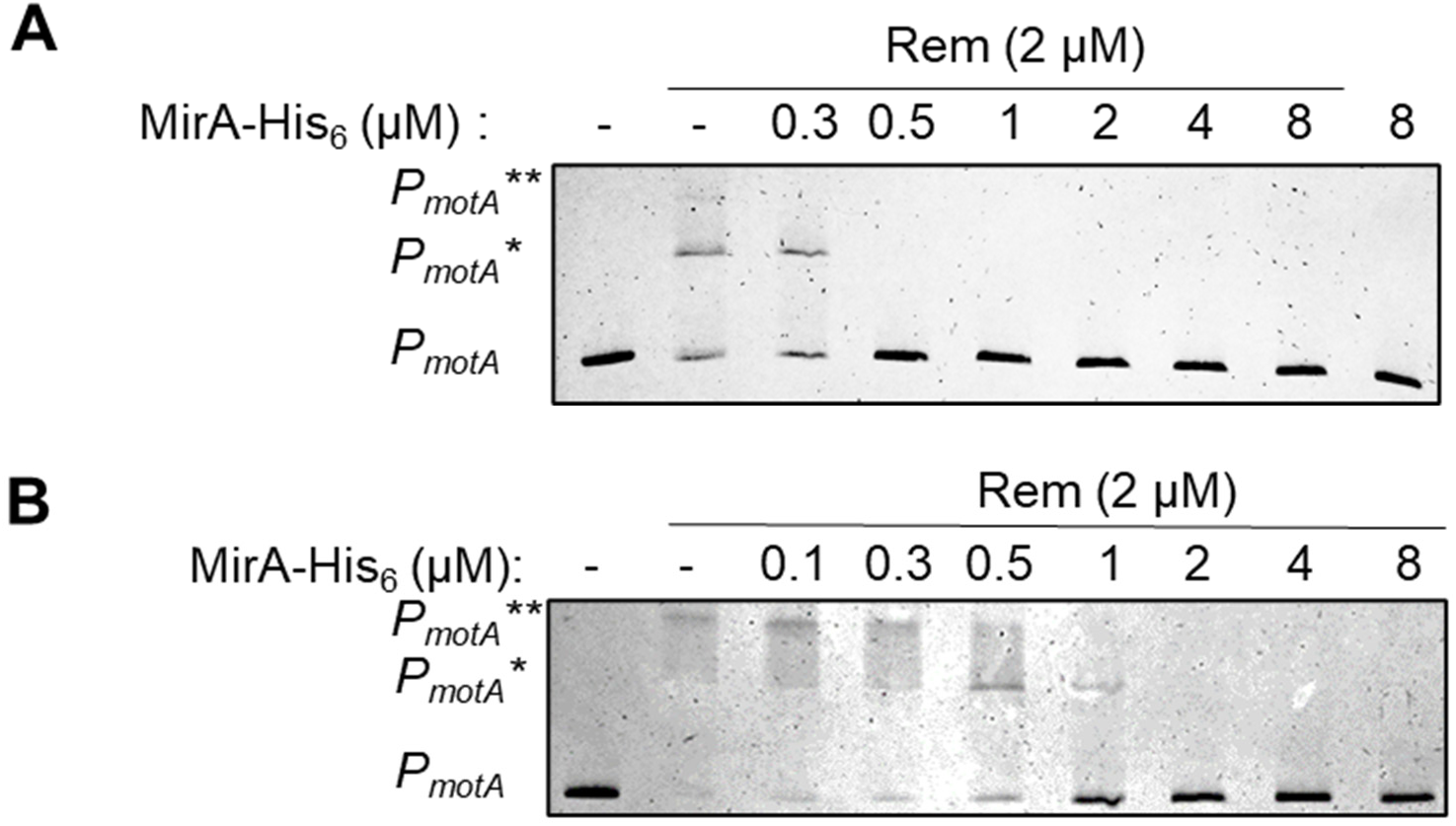
Gel shift inhibition of Rem by MirA. (A) Rem and MirA-His_6_ were pre-incubated 15 min followed by addition of 5 nM *P_motA_* DNA for a 15 min incubation all at RT. Reactions were separated on an 8% acrylamide gel and the DNA was stained with ethidium bromide (1 μg/mL) 15 min followed by imaging with UV by BioRad ChemiDoc. (B) Reactions were performed as in (A) but Rem was pre-incubated with 5 nM *P_motA_* DNA for 15 min prior, followed by the addition of MirA-His_6_.

### Conservation of MirA across the *Rhizobiaceae: A. tumefaciens* MirA inhibits motility in *S. meliloti*

The ExoR-ExoS-ChvI and the VisNR-Rem pathways have been well studied in *S. meliloti* (Chen *et al*., 2009, Rotter *et al*., 2006, Sourjik & Schmitt, 1996, Wells *et al*., 2007). We asked whether MirA functions similarly in *S. meliloti.* The *A. tumefaciens mirA* (*mirA*_At_) MirA was expressed in *S. meliloti* and the *S. meliloti mirA* (SM_17530, Smb20083, *mirA_Sm_)* ortholog in both *A. tumefaciens* and *S. meliloti* (Fig. 7A). Ectopic expression of *mirA_At_* inhibited motility in both species, but *mirA_Sm_* did not strongly alter motility in either species. We hypothesized that the strain of *S. meliloti* tested may have acquired mutations in *mirA* that decrease the protein’s interactions with both Rem*_Sm_* (SM_RS03450, Smc03046) and Rem*_At_*. To test whether MirA*_Sm_* interacts with Rem, we performed BACTH assays (Fig. 7B). The BACTH results suggested MirA*_Sm_* interacts with itself as well as MirA*_At_.* Weak interactions were observed between MirA*_Sm_* and Rem*_Sm_* and Rem*_At_.* Strong interactions were observed with Rem*_Sm_* and Rem*_Sm_* and Rem*_At,_* as well as MirA*_At_* with Rem*_Sm_.* We hypothesized that the Rem proteins may have a stronger sequence and structural conservation between *A. tumefaciens* and *S. meliloti* than the MirA homologs. Sequence alignments of homologs from different *Rhizobiales* species revealed that indeed, Rem proteins are highly similar across the *Rhizobiales* order (Figure S1). Significantly greater sequence divergence is observed when comparing MirA proteins from different species (indicated by asterisks)(Fig. S15). Since MirA from *A. tumefaciens* binds MirA*_At_* and MirA*_Sm_* and inhibits motility in both species, we hypothesized that the Rem proteins are similar enough that they would be able to cross-complement the *rem* mutants of each species. Indeed, introduction of *rem_Sm_* into the *A. tumefaciens* Δ*rem* and *rem_At_* into *S. meliloti* Δ*rem* complemented the motility defect of the *rem* deletion for both species, indicating that the *rem* genes can cross-complement (Fig. S16).

**Figure 7.**
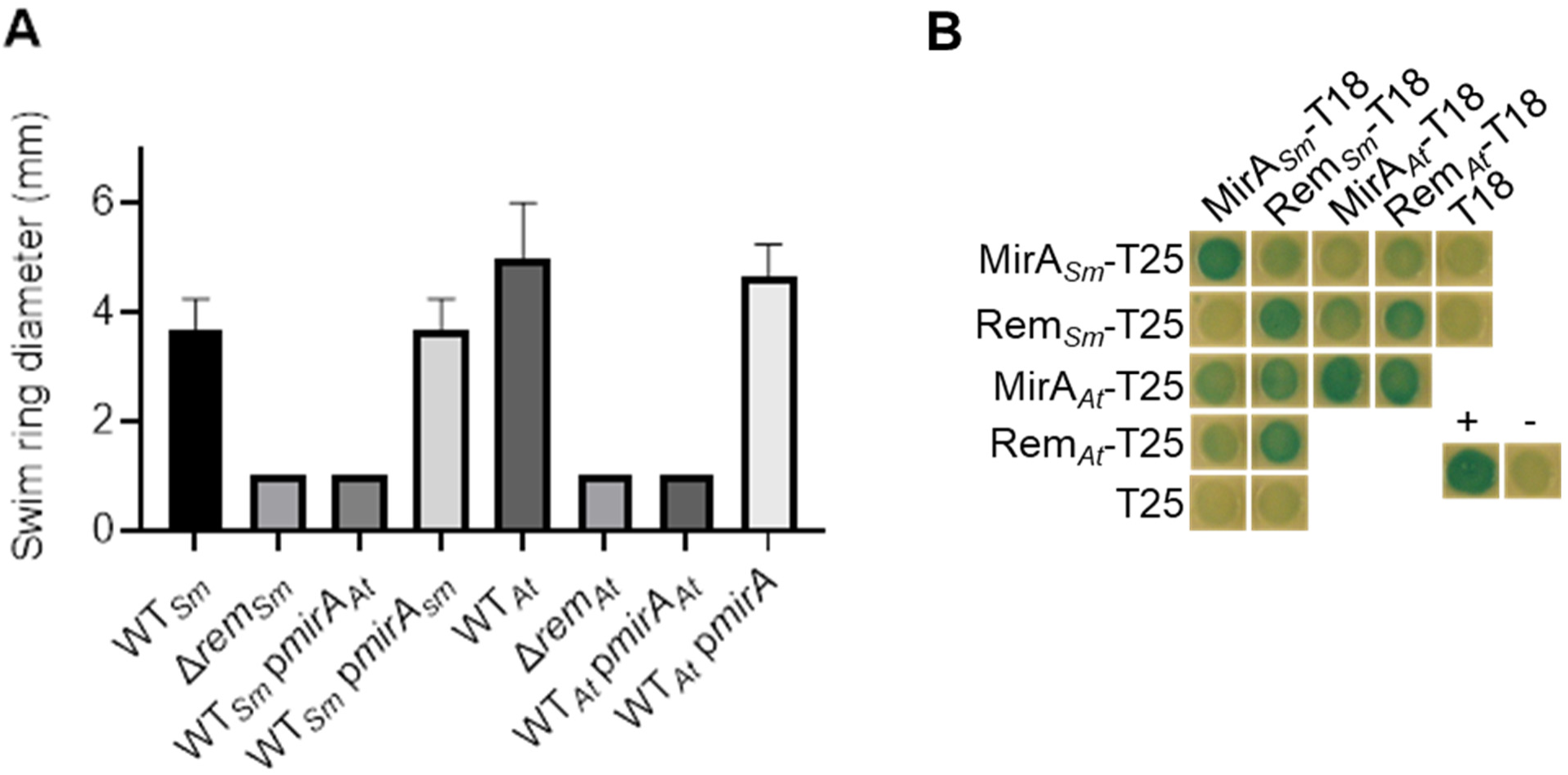
Ectopic expression of *A. tumefaciens* MirA can inhibit *S. meliloti* motility. (A) Swim motility assay of *A. tumefaciens* and *S. meliloti* expressing either *A. tumefaciens mirA_At_ or S. meliloti mirA_Sm_.* Colonies were inoculated into the center of 0.3% Bacto Agar (ATGN agar for *A. tumefaciens* and Bromfield agar for *S. meliloti*) and incubated 24 h in a humid chamber at RT. Error bars represent standard deviation for at at least 3 replicates (B) BACTH assay of MirA and Rem from *S. meliloti*. The *mirA*_Sm_ and *rem*_Sm_ coding sequences from *S. meliloti* strain 1021 were fused to the T25 and T18 domains of *E. coli* adenylate cyclase for BACTH and co-expressed in *E. coli.* Resulting *lacZ* activity was evaluated by plating 2 μL LB agar with X gal (80 μg/mL) and IPTG (500 μM) following a 24 h incubation.

## Discussion

In this study, we investigated the mechanism by which the ExoR-ChvG-ChvI pathway inhibits flagellar motility in *A. tumefaciens.* We identified a new regulator of flagellar gene expression in *A. tumefaciens,* a previously unannotated gene that encodes a 76-aa protein we have designated MirA. We find that MirA strongly inhibits motility gene expression (Fig. 4, Table S1), but that it does not affect transcription of the motility master regulators *visN*, *visR*, and *rem*. Instead, MirA forms a complex with the DNA-binding domain of the response regulator Rem, and this interaction inhibits the association of Rem with DNA (Fig. 5-6). Transcriptome data indicate that elevated *mirA* levels decrease transcription of the motility genes which are activated by Rem. We believe the different methods employed in our previous studies sufficiently explain the few MirA-regulated genes that were not identified as part of the VisR regulon (and therefore also the Rem regulon) by microarray (Heckel *et al*., 2014) (Table S2). Our data suggest that the primary function of MirA is to inhibit expression of motility genes in *A*. *tumefaciens* and that the ExoR-ChvG-ChvI regulators can direct this inhibition through activation of *mirA* expression (Fig. 8).

**Figure 8.**
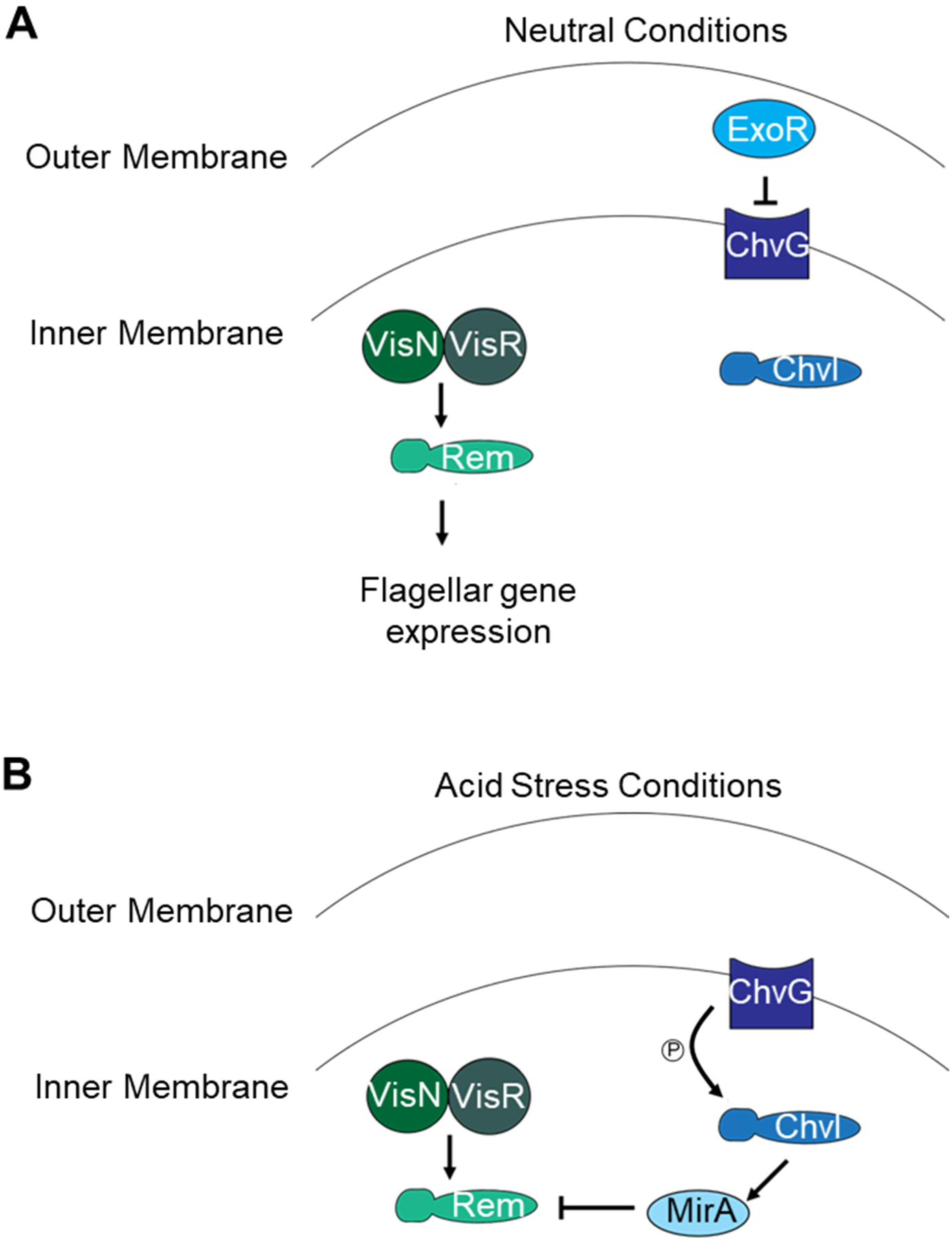
Model of MirA-dependent motility inhibition. (A) At neutral pH, VisN and VisR activate *rem* expression (Rotter *et al*., 2006, Xu *et al*., 2013). Rem directly activates expression of the class II motility genes, and flagella are expressed. ExoR inhibits activity of the ChvG-ChvI system (Wu *et al*., 2012) (B) In acidic pH, levels of ExoR protein decrease, de-repressing ChvG-Chvi protein, which leads to activation of *mirA* expression. MirA forms a complex with Rem, preventing it from binding to target promoters and thereby blocking motility gene expression.

### A new control mechanism for motility control in *A. tumefaciens*

Bacteria tightly regulate flagellar motility gene expression, only expressing genes for the components required as the flagellar nanomachine is being assembled. Flagellar gene expression is often regulated at multiple levels by a complex regulatory network of proteins and sRNAs (Osterman *et al*., 2015). A well-characterized inhibitory control mechanism is through the activity of anti-sigma factors, such as FlgM, that bind to and block the activity of motility-specific sigma factors required for transcription of late stage promoters including those that control expression of flagellin genes (Calvo & Kearns, 2015, Hughes *et al*., 1993). FlgM is secreted through the fully assembled type III secretion system flagellar nanomachine, releasing the cytoplasmic sigma factor to direct late-stage flagellar gene expression, thereby coupling biogenesis of the flagellar basal body to production of the flagellar filament. Although no specific sigma factor has been identified in the agrobacteria or rhizobia, the promoter elements for the class II flagellum biosynthetic genes suggest such an alternate sigma factor exists (Rotter *et al*., 2006). Initially, we hypothesized that MirA might function as an anti-sigma factor, but our results clearly show that MirA blocks flagellar assembly prior to class II gene expression through interaction with the Rem transcription factor.

We propose to add MirA to a small but growing list of post-transcriptional negative regulators of flagellar gene master regulators. In *Salmonella enterica,* YdiV is a regulatory protein that physically interacts with the FlhD flagellar activator protein, and can effectively remove the FlhD_4_C_2_ motility master regulatory complex from its target DNA in a similar fashion to how MirA can interfere with Rem and its ability to activate target promoters (Wada *et al*., 2011, Takaya *et al*., 2012). In contrast to MirA, YdiV is a much larger protein (237 aa) with a separate EAL domain found in phosphodiesterases that typically degrade c-di-GMP, although YdiV does not bind or turnover this second messenger (Simm *et al*., 2009). Interestingly, despite its larger size, YdiV acts to target the FlhD_4_C_2_ complex for proteolysis by ClpXP, in addition to blocking FlhD_4_C_2_ from binding to its target promoters. We cannot exclude the possibility that MirA functions as an adaptor for proteolysis of Rem in addition to inhibiting its DNA binding activity. Another post-transcriptional, negative regulator of the FlhD_4_C_2_ complex in *S. enterica*, FliT, interacts with FlhC, inhibiting motility by sequestering this regulatory component (Imada *et al*., 2010, Yamamoto & Kutsukake, 2006). Similar to MirA, FliT self-interacts (Imada *et al*., 2010). FliT acts as a secretion chaperone through interactions with filament cap protein FliD and interacts with the chaperone export protein FliJ (Evans *et al*., 2006). We have not observed any additional roles for MirA beyond interaction with the *A. tumefaciens* motility regulator Rem, but additional binding partners of MirA are possible.

In the aquatic APB *Caulobacter crescentus,* biogenesis of late-stage components of the polar flagellum is regulated by the MadA-FliX-FlbD system (Siwach *et al*., 2021). FliX is a 144-aa trans-acting regulator of flagellar gene transcription, and directly interacts with the FlbD motility regulator (Xu *et al*., 2011). FliX also associates with a gating component of the flagellar export machinery, FlhA, blocking export of late-stage flagellar substrates including the flagellins. The 91-aa protein MadA binds to the export machinery and stimulates displacement of FliX from FlhA, releasing it to interact with FlbD (Siwach *et al*., 2021). Interactions with FliX are required for FlbD to function, but at elevated levels also inhibit its DNA binding activity (Xu *et al*., 2011). In contrast, our results indicate that MirA is not required for Rem activity, but rather acts to strictly inhibit its ability to bind DNA. Additionally, MirA blocks expression of all the motility genes downstream of VisNR and Rem, rather than specifically affecting late-stage genes.

Our findings suggest that MirA functions an anti-activator of Rem. Beyond motility control, there are multiple examples of negative regulators, anti-repressors and anti-activators that directly interact with DNA-binding transcription factors to modulate their activities. In *A. tumefaciens,* the TraR quorum sensing transcriptional regulator is inhibited through direct interactions with its anti-activator TraM (Swiderska *et al*., 2001). TraM forms higher-order complexes with TraR dimers, blocking their ability to bind DNA and targeting TraR for proteolysis (Chen *et al*., 2007, Chen *et al*., 2004, Costa *et al*., 2012). Our BACTH results provide evidence that Rem and MirA both self-multimerize, but it is not yet clear whether these putative multimers dissociate during formation of the MirA-Rem complex, or rather assemble into higher-order complexes such as observed for TraM-TraR.

### Convergence of the ExoR-ChvG-ChvI and VisNR-Rem pathways

In our efforts to define the inhibitory mechanism of ExoR-ChvG-ChvI on motility, we gained insight into Rem and ChvI-dependent gene regulation in *A. tumefaciens*. Rem and its orthologs in other *Rhizobiales* have glutamate residues at the position that typically contains a phospho-accepting aspartate residue in two-component-type response regulators. It seemed plausible that the glutamate residue was acting as a stable phoshomimic, similar to what has been observed for other two-component response regulators (Klose *et al*., 1993). Mutation of this glutamate residue to asparagine did not abrogate Rem activity, nor did mutation of another aspartate proximal to this site (D45, Fig. S5). Similarly, substitution of single aspartate residues proximal to the glutamate residue at the predicted phosphorylation site of Rem*_Sm_* does not abolish Rem activity (Rotter *et al*., 2006). Cumulatively, these results suggest that Rem activity may not be regulated by phosphorylation, that the protein may be constitutively active, and this activity does not require the glutamate residue at position 50. Additionally, we found that, unlike other aspartate-less receiver domain-containing response regulators such as RitR (Glanville *et al*., 2018, Maule *et al*., 2015), Rem does not require the cysteine at amino acid 143, as replacement of the cysteine residue with serine, aspartate, or alanine did not impact Rem function (Fig. S6).

Mutation of the ChvI predicted phosphorylation site, however, did alter ChvI function *in vivo*. Mutation of the conserved aspartate residue at this position to asparagine (D52N) resulted in a loss-of-function allele, whereas mutation to a glutamate residue (D52E) acted as a phospho-mimic allele (Fig. 1), similar to other response regulators (Klose, 1993 #2031). Consistent with our expectations for a phospho-mimic ChvI allele, purified His_6-_ChvID52E binds to a target promoter (*P_hcpA_*) DNA *in vitro* (Wu *et al*., 2012). Unexpectedly, we found that purified wild-type His_6_-ChvI, which we assumed to be un-phosphorylated, was also able to bind P*_hcp_* in our assays (Fig. 2B). DNA binding *in vitro* by non-phosphorylated response regulators is not unheard of for two-component systems. In *S. meliloti,* the *chvID52E* allele has been shown to elevate succinoglycan production as detected by Calcofluor white fluorescence (Chen *et al*., 2009), which we also observed for *A. tumefaciens* (Fig. S2).

The ExoR-ChvG-ChvI system is a pervasive regulatory module in the *Rhizobiales* and has been proposed to regulate the free-living to host invasion switch (Heavner *et al*., 2015). MirA may represent a fourth component of this module, facilitating the motility regulation during this transition. Our results indicate that *mirA* expression is elevated in mutants expressing the *chvID52E* phosphomimic or in the *ΔexoR* mutant. Previous analysis of the Δ*exoR* transcriptome did not identify *mirA* as an ExoR-regulated gene (Heckel *et al*., 2014), but *mirA* was not yet annotated in the *A. tumefaciens* C58 reference genome at that time. Expression of *mirA* is elevated in acidic conditions. Although the ExoR-ChvG-ChvI pathway is recognized to enable response to acidic pH in *A. tumefaciens* and its relatives, there is growing evidence that this pathway may provide a more general stress response, and thus MirA-dependent control of motility may be relevant under a wider set of conditions than strictly acidic pH.

Multiple lines of evidence we present here indicate that in *A. tumefaciens* ChvI does not regulate *visN, visR,* or *rem* gene expression, nor does it bind directly to Rem nor the promoters of motility genes (Fig. 1B, Fig. S3, Fig. 2D, Fig. S8). Rather, ChvI inhibits motility through activating expression of *mirA*. In *S. meliloti*, ChvI has been reported to directly regulate *rem* expression, in contrast to our findings in *A. tumefaciens* (Heckel *et al*., 2014, Wells *et al*., 2007, Ratib *et al*., 2018). Ectopic expression of the *S. meliloti mirA* (SM_RS17530, Smb20083, MirA_Sm_) homolog did not inhibit motility in either *S. meliloti* or *A. tumefaciens* (Fig. 7A), whereas ectopic expression of *mirA*_At_ inhibited motility in both bacterial taxa. The inability of MirA_Sm_ to inhibit motility may be reflective of its high sequence divergence relative to other *mirA* homologs (Fig. S15) and is consistent with its weak interaction with Rem from either *S. meliloti* or *A. tumefaciens* (Fig. 7C). It is interesting that control of motility by ExoR-ChvG-ChvI has been conserved between *A. tumefaciens* and *S. meliloti*, although the endogenous regulatory mechanism employed seems to have diverged. We observed self-interaction of both MirA*_Sm_* and Rem*_Sm_*. The ability of MirA_At_ to inhibit *S. meliloti* motility is predicted based on the high level of sequence conservation between the Rem_At_ and Rem_Sm_ proteins (Fig. 7b, Fig. S1). There are putative, better conserved MirA homologs in multiple other *Rhizobiales* species which also encode the Rem transcription factor (Fig. S1), suggesting that similar mechanisms of MirA-directed motility control exist in these systems.

## Experimental Procedures

### Bacterial strains, plasmids, and oligonucleotides

Bacterial strains and plasmids are listed in Table S3, and oligonucleotides are listed in Table S4. *E. coli* strains were grown in LB at 37 ⁰C and *A. tumefaciens* strains were grown in LB and AT minimal medium with 15 mM (NH_4_)_2_SO_4_, buffered to pH 7 with KH_2_PO_4_ to a final concentration of 79 mM with 0.5% glucose as the carbon source (ATGN). Acidic medium was prepared with a final concentration of 20 mM MES (2-(*N*-morpholino)ethanesulfonic acid) hydrate buffered to pH 5.5 in place of KH_2_PO_4_ buffer. *S. meliloti* strains were grown in YTC media (0.5% tryptone, 0.3% yeast extract, .08% CaCl_2_· 2H_2_0 pH 7.0). Growth temperatures were 28-30 ⁰C for *S. meliloti* and *A. tumefaciens* and 37 ⁰C for *E. coli*. Antibiotics were used at the following concentrations for liquid media: Kanamycin (Km) 150 μg/mL, Gentamycin (Gm) 100 μg/mL, Carbenicillim (Cb) 25 μg/mL, Spectinomycin (Sp) 150 μg/mL for *A. tumefaciens*; Km 25 μg/mL, Gm 25 μg/mL, Ampicillin (Ap) 100 μg/mL, Sp 25 μg/mL for *E. coli*; Gm 40 μg/mL for *S. meliloti*.

### Allelic replacement and marker-less deletion construction

Marker-less deletion strains was performed as previously described (Morton & Fuqua, 2012). Fragments upstream and downstream of the genes of interest (0.5-1 kb in size) were amplified using primers P1 and P2 (upstream) and P3 and P4 (downstream) (Table 2). These fragments were stitched together via 18-20 base-pair complementary sequences by a SOEing reaction. The deletion or allelic replacement construct was first ligated into the pGEM T-Easy vector for amplification in *E. coli* strain Top 10 F.’ Mini-prepped pGEM plasmids were digested with the appropriate enzymes and the insert ligated into the suicide vector pNPTS138 (Hibbing & Fuqua, 2011). This vector was transformed into the *E. coli* conjugal donor strain S17-1/λ*pir* to transfer the deletion plasmid into the appropriate *A. tumefaciens* strain. Mixtures of *A. tumefaciens* recipient and *E. coli* donor strains were spotted onto cellulose acetate filter disks as described previously (Morton & Fuqua, 2012). Primary integrants were selected by plating cells collected from filter disks onto ATGN-Km plates. Cells that subsequently lost the integrated plasmid but retained the introduced deletion were isolated through passage in the absence of antibiotic by plating onto AT medium with 5% sucrose instead of glucose (ATSN) and confirmed by PCR across the deletion junction.

### Construction of expression plasmids

Controlled expression plasmids were generated by introducing *A. tumefaciens* coding sequences amplified from genomic DNA by PCR into the LacI^Q^-encoding, IPTG (isopropyl-β-D-thiogalactopyranoside)-inducible expression vector with a gentamicin resistance (Gm^R^) cassette, pSRKGm (Khan *et al*., 2008). For expression *mirA* from its native promoter and *P_lac_,* the *mirA* coding sequence and upstream region were cloned into pSRKGm at the *Nde*I site, with a stop codon to prevent translation across the *Nde*I site from the upstream *lacZα*. To construct *P_mirA_*::*mirA_Sm_*, the *mirA* gene from *A. tumefaciens* was replaced by the *mirA* gene from *S. meliloti* on pSRK:: *P_mirA_*::*mirA_At_*. For expression from the phage promoter *P*_N25_, *mirA* was cloned into pYW15c (Wang *et al*., 2000). Coding sequences were amplified from *A. tumefaciens* C58 genomic DNA or plasmids carrying engineered alleles using the 5’ and 3’ primers corresponding to each gene, specified in Table S4. Amplified fragments were ligated into the cloning vector pGEM-T Easy (Promega), or built directly into the plasmid of interest using the NEBuilder HiFi Assembly kit (New England Biolabs), transformed into *E. coli* Top10 F’ or DH5α/λpir for amplification and collection, and the inserts were confirmed by sequencing. Coding sequences were excised from the pGEM-T Easy cloning vectors by restriction enzyme cleavage and ligated into the appropriately cleaved pSRKGm vector using T4 DNA ligase (New England Biolabs). Derived plasmids were verified by PCR amplification across the multiple cloning site and insert prior to transformation into *A. tumefaciens* cells via electroporation (Mersereau *et al*., 1990).

### Site-directed mutagenesis of ChvI and Rem

Site-directed mutagenesis was performed to create the alleles *chvID52E*, *chvID52N, remD45N*, *remE50N, remC143S, remC143D,* and *remC143A*. Mutagenesis was performed as previously described (Hibbing & Fuqua, 2011, Mohari *et al*., 2018) using the protocol described in the QuickChange Site-Directed Mutagenesis Kit (Stratagene Corp) or the Q5 Site-Directed Mutagenesis Kit (New England Biolabs). The desired nucleotide changes were designed into the complementary mutagenesis primers (Table S4). Mutant alleles were amplified from cloning constructs carrying the wild-type alleles. After amplification of the entire plasmid, template and hemi-methylated plasmid DNA were removed from the reaction via *DpnI* digestion. The mutagenized plasmids were transformed into *E. coli* Top10 F’ or DH5α/λpir, collected, sequence verified, and the alleles were fused with controlled expression plasmids or allelic replacement plasmids.

### Construction of *lacZ* fusions and β-galactosidase assays

Promoter fusions to *lacZ* were constructed by amplification of promoter regions from genomic DNA with the specified primers (Table S4) and ligated into pGEM-T Easy (Promega) for amplification and collection. Promoter fragments were digested from cloning constructs and ligated into reporter plasmid pRA301 (Akakura & Winans, 2002) to generate reporter constructs with the promoter region, ribosome binding site, and start codon of the gene of interest fused in-frame with the second codon of the *lacZ* gene to generate translational fusions.

Bacterial cultures were grown to exponential phase at 30 ⁰C and frozen in 30% glycerol at OD_600_ 12. Thawed samples were inoculated at OD_600_ 0.05 in 2 mL cultures in triplicate, grown to exponential phase at 30 ⁰C, measured for OD_600_ and frozen. Thawed samples were incubated in Z buffer (Miller, 1972) with 2 drops of 0.02% SDS, 4 drops of chloroform, vortexed for 10 seconds, and incubated with 100 μL 4 mg/mL ortho-nitrophenyl-β-D-galactopyranoside (ONPG) at RT. Timed reactions were stopped with addition of 600 μL NaCO_3_ and clarified through centrifugation. β-galactosidase cleavage of ONPG was measured through A_420_ measurement in a BioTek plate reader and specific activity was measured in Miller Units (Miller, 1972).

### 5’RACE mapping transcriptional start sites

Total RNA was extracted from either wild-type *A. tumefaciens* (for mapping the TSS of *motA* and *flgB*) or Δ*exoA*Δ*exoR* (for mapping the TSS of *hcp, chvI,* and *aopB*). RNA was extracted using the RNeasy Midi Kit from QIAGEN (Valencia, CA). Residual DNA was removed from the extracted RNA using the TURBODNA-*free*_TM_ Kit from Thermo Fisher Scientific (formerly Life Technologies). Enzymes provided in the TAKARA 5’-Full RACE Core Set were used for 1_st_ strand cDNA synthesis, degradation of hybrid DNA-RNA, and circularization of single-strand cDNA, according to the kit instructions. Synthesis of cDNA was directed by 5’-phosphorylated primers (Table S4). Amplification of promoter regions was performed with Phusion polymerase (NEB) with the A1, S1, A2, and S2 primers (Table S4) according to the TAKARA 5’-Full RACE Core Set kit instructions. Amplified promoter fragments were ligated into the pGEM-T Easy (Promega) and transformed into *E. coli* Top10 F’ for collection and sequencing.

### RNA sequencing

1.5 mL exponential phase cultures were mixed with 1.5 mL RNAProtect Reagent. RNA was purified with the Qiagen RNEasy kit, and DNA was digested with the Turbo RNAse-free DNAse kit for 1 hr at 37 ⁰C. The integrity of the RNA was checked with Agilent Tapestation. From each sample, 1 ug of total RNA was used for ribosomal RNA depletion with the Ribo-Zero™ Magnetic Kit (Bacteria)(Epicenter). The libraries were then prepared with the TruSeq Stranded mRNA library preparation kit (Illumina). The cleaned adapter-ligated libraries were pooled and loaded on the NextSeq 500 with a 75 high cycle sequencing module to generate paired-end reads. Mutations were identified by mapping reads against the *A. tumefaciens* C58 reference genomes (Genbank accession numbers AE008687, AE008688, AE008689, and AE008690 for the circular, linear, At, and Ti plasmids, respectively) using the *breseq* computational pipeline (Deatherage & Barrick, 2014). Transcriptome data is summarized in Table S1 and is available at GEO-NCBI (GSE174467).

### Motility assays

To quantify the efficiency of flagellar locomotion, strains were inoculated into the center of motility medium plates with 0.3 % Bacto Agar suspended in media (ATGN for *A. tumefaciens*, Bromfield agar (Sourjik, 1996 #1979) for *S. meliloti*) and the diameter of each swim ring was measured over time. Strains were inoculated from single colonies from 1.5% agar (ATGN or 2xYT) plates with a toothpick or wire. Swim plates were incubated at RT in a sealed container with an open vial of a saturated K_2_SO_4_ solution to maintain a relative humidity of ∼97% in the chamber. Swim ring diameters were measured with a standard ruler.

### Biofilm assays

Biofilm assays were performed as previously described (Tomlinson *et al*., 2010). Cultures of *A. tumefaciens* were inoculated at an initial OD_600_ 0.05 in ATGN supplemented with 22 μM FeSO_4_ ∙ 7H_2_O into 12-well plates with upright polyvinyl chloride (PVC) coverslips or into 96-well PVC plates for large-scale assays. Plates were incubated at RT 24-48 h in a sealed container with an open vial of a saturated solution of K_2_SO_4_ to maintain a relative humidity of ∼97% in the chamber. The coverslips or wells were stained with 0.1% crystal violet dye and the dye was solubilized in 33% acetic acid. Biofilm biomass was determined by measuring the A_600_ of solubilized crystal violet and normalizing by the OD_600_ of the cultures at the time of harvesting. The cell density and absorbance readings were measured with a Bio-TEK Synergy HT plate reader using Bio-TEK Gen5 (version 1.07) software.

### Calcofluor white staining

ATGN plates were supplemented with 200 μg/mL of Calcofluor White (fluorescent white) dye from a stock solution of 20 mg/mL. *A. tumefaciens* strains were inoculated into ATGN liquid medium from colonies and grown overnight at 28°C with shaking. Cultures were normalized to an OD_600_ 0.2, and 5 μL of each culture was spotted onto the plate. Plates were incubated for 2 days at 28°C and then imaged with UV light excitation using a Biorad ChemiDoc system with Image Lab software. All strains were grown and imaged on the same plate.

### Isolation and characterization of motile suppressors of Δexo*R*

Motile suppressor mutants of Δ*exoR*Δ*exoA* were isolated as described previously (Heckel *et al*., 2014). Briefly, the nonmotile Δ*exoR*Δ*exoA* mutant strain was mutagenized with the *mariner* transposon (*Himar1*) and inoculated into 0.3% Bacto Agar dissolved in ATGN media. Motile suppressor mutants were harvested from the edge of the swim ring. To identify suppressors with mutations beyond the *chvG*-*chvI* locus, which restore both motility and biofilm formation to the Δ*exoR* mutant strain, suppressor mutants were screened for the ability to form biofilms. We used touchdown PCR and sequencing to identify the site of transposon insertion for these mutants as well as whole genome sequencing to identify point mutations in these genomes.

### Whole genome sequencing

Genomic DNA was prepared from 1 mL of cultures grown in ATGN medium to mid-log phase. Genomic DNA of the Δ*exoA*Δ*exoR* parent and four suppressor mutants was prepared and used to create libraries for sequencing. Libraries were prepared using a Bio Scientific NEXTflex™ Rapid DNA sequencing kit according to manufacturer instructions. Sequencing was performed with an Illumina Nextseq instrument with subsequent analysis from the IU CGB. Mutations were identified by mapping reads against the *A. tumefaciens* C58 reference genomes (Genbank accession numbers AE008687, AE008688, AE008689, and AE008690 for the circular, linear, At, and Ti plasmids, respectively) using the *breseq* computational pipeline (Deatherage & Barrick, 2014).

### Expression and purification of proteins

The *rem* allele was amplified from a cloning construct (primer sequences in Table S4) and ligated into pTYB12 (NEB IMPACT system of protein purification), resulting in an N-terminal intein fusion to Rem. The expression plasmid was transformed into *E. coli* Top10 F’ for sequence verification and collection and then transformed into *E. coli* BL21/λDE3 for expression and purification. Expression and purification of the intein-tagged Rem protein was performed according to the protocol described in the NEB IMPACT_TM_ Kit. One liter of cells was grown at 37°C on an orbital shaker to an OD_600_ 0.5 then induced with IPTG to a final concentration of 400 μM. Cells were induced overnight on an orbital shaker at 16°C. Cells were collected by centrifugation at 5,000 x g for 15 min at 4°C, and the pellet was re-suspended in column buffer (20 mM Tris-HCl pH 8.5, 500 mM NaCl, 1 mM EDTA). Cells were lysed by passage through an M-110L Microfluidizer Processor (Microfluidics, Westwood,MA). Cell debris was removed by centrifugation at 15,000 x g for 30 min at 4°C and the clarified extract was loaded onto a chitin resin (NEB) column pre-equilibrated with column buffer for purification at RT. After washing with 20 bed volumes of column buffer, auto-cleavage of the intein tag was induced by adding 3 bed volumes of cleavage buffer (column buffer with 50 mM DTT) to the column and incubating at RT for 40 h. Un-tagged Rem protein was eluted with column buffer and pooled fractions were dialyzed into a storage buffer (10 mM Tris-HCl, pH 8.0, 250 mM NaCl, 1 mM DTT, 10%glycerol). Aliquots were flash frozen and stored at −80°C.

The *chvI* and *mirA* coding sequences (wild type and mutant alleles) were amplified from cloning constructs (primer sequences in Table 2) and ligated into pET-28a(+) (Novagen/ EMB Biosciences/ Millipore). Protein-expression plasmids were transformed into *E.coli* Top10 F’ for collection and sequence verification and then transformed into *E.coli* BL21/λDE3 for expression and protein purification. One liter of cells were grown in LB at 37°C on an orbital shaker to an OD_600_ of 0.6, then induced with 500 μM IPTG for 4 hours at 37°C. Cells were collected by centrifugation at 4°C at 5,000 x g for 15 min. Cell pellets containing ChvI were resuspended in lysis buffer (50 mM NaH_2_PO_4_ pH 6.5, 300mM NaCl, 10 mM imidazole) with 1 mM PMSF (phenylmethanesulfonyl fluoride). Cells were lysed by passage through an M-110L Microfluidizer Processor (Microfluidics). Cell debris was removed by centrifugation at 15,000 x g for 30 min at 4°C and pellets were resuspended in storage buffer (50 mM NaPO_4_ pH 6.5, 500 mM NaCl). Purification was performed on 1 ml HisTrap HP (GE Healthcare) column using an FPLC system (ÄKTA P-920 pump and Superdex 75 10/300 GL column, GE Healthcare). Protein-containing fractions were pooled for desalting through a 5 mL HiTrap Desalting column (GE Healthcare). Desalted protein was dialyzed into a storage buffer (50 mM NaH_2_PO_4_ pH 8.0, 500 mM NaCl, 20% glycerol). Protein concentration was calculated following absorbance measurement by Nanodrop with the molar extinction coefficient for each protein.

Cell pellets from cells expressing MirA-His_6_ were thawed and suspended in 20 mM NaH_2_PO_4_, 150 mM NaCl, pH 7.4 buffer with 1 mM PMSF and cells were disrupted by passage through an EmulsiFlex-C3 emulsifier, and clarified by centrifugation. Clarified lysates were loaded onto a 1 mL TALON (Takara) column, washed with 20 mM NaH_2_PO_4_, 150 mM NaCl, pH 7.4 buffer and eluted with 20 mM NaH_2_PO_4_, 150 mM NaCl, 5% glycerol, pH 7.4 buffer with 100–600 mM imidazole step elution fractions. Eluted fractions were pooled and dialyzed into 20 mM NaH_2_PO_4_, 150 mM NaCl, 5% glycerol, pH 7.4 buffer and frozen at −20 °C. Protein concentration was determined by Pierce BCA assay as molar absorptivity measurements with MirA-His_6_ were clearly erroneous.

### Electrophoretic mobility shift assay

Upstream DNA fragments suspected to have promoter elements were amplified and purified with the E.Z.N.A. Cycle Pure kit (Omega Bio-Tek). 10 nM DNA fragments were incubated with purified protein in binding buffer (10 mM Tris-Cl pH 7.5, 1 mM EDTA, 0.1 mM DTT, 5% glycerol, and 0.05 mg/mL BSA) and incubated 20 min at RT. Reactions were run on 6% acrylamide (80:1 acrylamide: bisacrylamide) 0.5% TBE (45 mM Tris-borate, 1 mM EDTA pH 8.0) gels and electrophoresed for 2 h at 150 V. Prior to loading, gels were pre-run with 0.5% TBE at 75 V for 1 h. Gels were stained with a 10,000:1 dilution of SYBR Safe DNA gel stain (Invitrogen) 20 min in 0.5X TBE, rinsed with distilled water and imaged on a Bio-Rad ChemiDoc system using Image Lab software. For gel shift inhibition assays, purified MirA-His_6_ protein was incubated with Rem protein for fixed incubation times at RT prior to the addition of DNA, followed by 15 min incubation at RT. Reactions were run on an 8% polyacrylamide gel prior to staining for 15 min with 1 μg/mL ethidium bromide and imaging on a Bio-Rad ChemiDoc.

### Pull-down assay

Purified proteins were diluted to the appropriate concentration in 50 mM Tris-HCl, 100 mM NaCl, 0.1% Tween pH7.5 buffer. Samples were incubated 30 min at RT and loaded onto a 10 μL aliquot of Talon (Takara) resin. Following a 30 min incubation, the resin was washed 3 times with 500 μL buffer and eluted with 20 μL Laemlli loading buffer.

### Western and farwestern blotting

For farwestern analysis, purified protein was electrophoresed by SDS-PAGE in two identical gels (12.5% acrylamide, 37.5:1 acrylamide:bisacrylamide) and transferred to nitrocellulose membranes. Membranes were blocked with 5% nonfat dry milk in TBS 0.1% Tween for 1 hour. For MirA-_6_His, membranes were blotted either with milk containing purified 0.4 M MirA-His_6_ or buffer. For _6_His-ChvI^D52E^, the proteins on the membrane were denatured by incubation in AC buffer (100 mM NaCl, 20 mM Tris, 0.5 mM EDTA, 10% glycerol, 0.1% Tween-20) with 6 M guanidine-HCl for 30 min at RT with mixing. Proteins were renatured by incubating at RT with shaking for 30 min in AC buffer with 3M guanidine-HCl, then 30 min at RT in AC buffer with 1 M guanidine-HCl, then 30 minutes at 4°C in AC buffer with 0.1 M guanidine-HCl, then overnight at 4°C in AC buffer. The membranes were blocked in blotto (%5 milk solution in tris-buffered saline with 1% Tween-20) at RT 1 h with shaking. Purified His_6_-ChvID^52E^ protein (5 μg) was used as prey and incubated with a membrane in protein binding buffer (AC buffer with 5% milk powder and 1 mM DTT) overnight at 4°C with shaking. Another membrane was processed in tandem in the same conditions with no His_6_-ChvI^D52E^. Membranes for both proteins were then incubated with a 1:2000 dilution of rabbit α-His antibody for 1 hour at RT, then with a 1:20,000 dilution of HRP-conjugated goat anti-rabbit antibody for 1 hour at RT. For detection of MirA-FLAG_3_, strains were grown to exponential phase, concentrated to an OD_600_ of 10, and boiled 5 min prior to running on a 12.5% acrylamide (37.5:1 acrylamide:bisacrylamide) SDS-PAGE gel with 2,2,2 trichloroethanol (TCE). The TCE was UV-activated prior to transfer to a nitrocellulose membrane. Membrane transfer was confirmed through UV-imaging of membrane (BioRad ChemiDoc ImageLab software). The membranes were incubated with 1:5,000 monoclonal ANTI-FLAG M2 antibody produced in mouse (Sigma-Aldrich) and Goat Anti-Mouse IgG (H+L)∼HRP Conjugate (BioRad). Membranes were developed with SuperSignal® West Pico Luminol/Enhancer Solution (Thermo Scientific) and the HRP signal was detected using a Biorad ChemiDoc system using Image Lab software.

### Bacterial adenylate cyclase two-hybrid assays

Genes of interest were PCR-amplified and inserted into *Kpn*I and *Eco*RI-digested pKNT25 or pUT18 with NEB Hifi Assembly to construct in-frame C-terminal fusions (Euromedex). Plasmids containing T18 and T25 fusions were co-transformed into BACTH test strain BTH101. Strains were grown overnight at 30 ⁰C and 2 μL were spotted onto LB plates containing IPTG (500 μM) and 5-bromo-4-chloro-3-indolyl-β-D-galactopyranoside (X-gal, 80 - 160 μg/mL) and incubated at 30 ⁰C overnight. Colonies were imaged with a Nikon camera. All strains were grown and imaged on the same plate for each experiment.

### Multiple Sequence Alignment

Amino acid sequences were retrieved from Kegg, Biocyc or JGI IMG and aligned using ClustalW with the default parameters (Gap open penalty = 10, Gap extension penalty = 0.5, BLOSUM weight matrix). Amino acid sequences of proteins can be found at NCBI reference numbers WP_006313038.1 (Rem/*A. tumefaciens*), CAC45250.1 (Rem/*S. meliloti*), SUB44985.1 (Rem/*O. anthropi*), WP_006698188.1 (Rem/*R. pusense* IRGB74), WP_065114859.1 (Rem/*A. rhizogenes*) WP_010970615.1 (ChvI/*A. tumefaciens*), WP_003531999.1 (ChvI/*S. meliloti*), AYM61105.1 (VirG/*A. tumefaciens*), NP_417864.1 (OmpR/*E. coli*), AKH06072.1 (PhoB/*S. enterica*), WP_003502959.1 (MirA/ *A. tumefaciens*), CAC48483.1 (MirA/ *S. meliloti*), WP_004442019.1 (MirA/*R. pusense* IRGB74), WP_065115793.1 (MirA/*A. rhizogenes*), and WP_040128199.1 (MirA/*Ochrobactrum/Brucella anthropi*).

## Acknowledgements

This project was supported by National Institutes of Health (NIH) grant GM120337 (C.F.) Illumina DNA Sequencing and bioinformatic analysis of whole genome re--sequencing and RNAseq were performed by the Indiana University Center for Genomics and Bioinformatics.

## Author contributions

C.F. formulated the research plan; M.A.A, B.C.H., A.S., J.A.S. and C.F. performed the experiments, C.F., M.A.A. and B.C.H. wrote the paper.

## Data availability

High-throughput, Illumina-based RNASeq data has been deposited in the GEO database Accession No. GSE174467 (([dataset] Alakavuklar and Fuqua, 2021) (Deatherage et al, 2014)). All other data that support the findings of this study are available from the corresponding author upon reasonable request.

