## Supplemental Material for "Motility Control Through an Anti-Activation Mechanism in *Agrobacterium tumefaciens*"

Supplementary Tables: S3-S4 (S1 and S2 are large Excel files provided separately)

Supplementary References:

Supplementary Figure Legends:

Supplementary Figures: S1-S16

**Table S3. Strains and plasmids used in this study**

| Strain/plasmid | Relevant features | Reference |
|---|---|---|
| ***Escherichia coli*** | | |
| DH5α/λ*pir* | λ*pir*; cloning strain | (Woodcock *et al.*, 1989) |
| S17-1/λ*pir* | λ*pir*; $Tra^+$; cloning host | (Kalogeraki & Winans, 1997) |
| Top10F' | Cloning strain | Invitrogen |
| BL21/DE3 | Protein expression strain | (Tabor & Richardson, 1985) |
| ***Agrobacterium tumefaciens* C58 strains** | | |
| C58 | Nopaline-type; pTiC8, pAtC58 | (Watson *et al.*, 1975) |
| PMM1 | Δ*exoR* (ATU_RS08400, Atu1715) | (Heckel *et al.*, 2014) |
| PMM2 | Δ*exoA*Δ*exoR* (ATU_RS08400 ATU_RS18935, Atu4053, Atu1715) | (Tomlinson *et al.*, 2010) |
| BCH101 | Δ*exoR*Δ*chvI* (ATU_RS08400 ATU_RS00165 / Atu1715, Atu0034) | (Heckel *et al.*, 2014) |
| BCH103 | Δ*chvI* (ATU_RS0016 / Atu0034) | (Heckel *et al.*, 2014) |
| BCH122 | ChvID52N chromosomal allele (ATU_RS00165 / Atu0034) | This study |
| BCH123 | ChvID52E chromosomal allele (ATU_RS00165 / Atu0034) | This study |
| BCH124 | Δ*exoR*ChvID52N (ATU_RS08400, ATU_RS00165 / Atu1715, Atu0034) | This study |
| BCH125 | Δ*exoR*ChvID52E (ATU_RS08400, ATU_RS00165 / Atu1715, Atu0034) | This study |
| JX148 | Δ*rem* (ATU_RS02820 / Atu0573) | This study |
| BCH117 | Δ*exoR*Δ*rem* (ATU_RS08400, ATU_RS02820 / Atu1715, Atu0573) | This study |
| BCH116 | Δ*chvI*Δ*rem* (ATU_RS00165, ATU_RS02820 / Atu0034, Atu0573) | This study |
| BCH132 | Δ*exoA*Δ*exoR*, motility suppressor, Tn::*atu2241*, C→G at bp 1,625,562 on circ. chrom., $Km^R$ | This study |
| BCH133 | Δ*exoA*Δ*exoR*, motility suppressor, Tn::*atu2241*, ΔA at bp 1,625,665 on circ. chrom., $Km^R$ | This study |
| BCH134 | Δ*exoA*Δ*exoR*, motility suppressor, Tn::*atu2241*, C→A at bp 1,625,700 on circ. chrom., $Km^R$ | This study |
| BCH136 | BH133 mutation introduced to wt C58 | This study |
| BCH137 | BH133 mutation introduced to Δ*exoA*Δ*exoR* | This study |

| | | |
|---|---|---|
| MAT3 | Δ*exoA*Δ*exoRmirA*ATC (ATU_RS08400, ATU_RS18935, ATU_RS08050 / Atu4053, Atu1715) | This study |
| MAT4 | Δ*exoA*Δ*exoR*Δ*mirA (*ATU_RS08400, ATU_RS18935, ATU_RS08050 / Atu4053, Atu1715*)* | This study |
| MAT5 | Δ*mirA* (ATU_RS08050) | This study |
| MAT6 | Δ*exoA*Δ*exoR*Atu1638S20STOP (ATU_RS08400 ATU_RS18935, ATU_RS08050 / Atu4053, Atu1715) | This study |
| MAT7 | *ΔexoAΔexoRmirAG20*STOP (ATU_RS08400 ATU_RS18935, ATU_RS08050 / Atu4053, Atu1715) | This study |
| MAT8 | *mirA*-FLAG3 (ATU_RS08050) | This study |
| MAT9 | *ΔexoAΔexoRmirA*-FLAG3 (ATU_RS08400 ATU_RS18935, ATU_RS08050 / Atu4053, Atu1715) | This study |
| ***Sinorhizobium meliloti*** | | |
| RM1021-KEG2108 | $Sm^R$, $Cm^R$, pLAFR2070 (cbrA) Δ*cbrA* | (Gibson *et al.*, 2006) |
| Ru11/001 | $Sm^R$, spontaneous streptomycin-resistant isolate; wild-type strain | (Pleier & Schmitt, 1991) |
| Ru11/555 | $Sm^R$, Δ*rem* (Smc03046) | (Rotter *et al.*, 2006) |
| **Plasmids** | | |
| pGEM-T Easy | Cloning vector, $Amp^R$ | Promega |
| pNPTS128 | ColE1 suicide plasmid; *sacB*; $Km^R$ | Gift of M. Alley |
| pET-28a (+) | His tag fusion vector, $Km^R$ | Novagen |
| pTYB12 | N-terminal Intein tag fusion vector, $Amp^R$ | New England Biolabs |
| pSRKGm | Broad host range $P_{lax}$ expression vector; *lacIQ*; $Gm^R$ | (Khan *et al.*, 2008) |
| pRA301 | Broad host range; promoterless *lacZ*; $Spec^R$ | (Akakura & Winans, 2002) |
| pKNT25 | T25 fragment fused downstream of MCS; pSU40 derivative, $Km^R$ | Euromedex |
| pUT18 | T18 fragment fused downstream of MCS, pUC19 derivative; $Amp^R$ | Euromedex |
| pKT25-zip | T25-Zip fusion, positive control for BACTH assay, $Km^R$ | Euromedex |
| pUT18C-zip | T18-Zip fusion, positive control for BACTH assay, $Amp^R$ | Euromedex |
| pYW15c | Broad host range $P_{N25}$ vector; $Amp^R$ | (Wang *et al.*, 2000) |
| pJX160 | pRA301::*PflgE* (ATU_RS02825 / Atu0574)-*lacZ*; $Spec^R$ | (Xu *et al.*, 2013) |
| pJX166 | pRA301::*PmotA* (ATU_RS02760 / Atu0560)-*lacZ*; $Spec^R$ | (Xu *et al.*, 2013) |

| | | |
|---|---|---|
| pJX159 | pRA301::*Prem* (ATU_RS02820 / Atu0573)-*lacZ;* $Spec^R$ | (Xu *et al.*, 2013) |
| pJX145 | pSRKGm::*rem* (ATU_RS02820 / Atu0573); $Gm^R$ | This study |
| pBCH155 | pSRKGm::RemD45N (ATU_RS02820 / Atu0573); $Gm^R$ | This study |
| pBCH156 | pSRKGm::RemE50N (ATU_RS02820 / Atu0573); $Gm^R$ | This study |
| pBCH111 | pRA301::$P_{aopB}$-*lacZ* (ATU_RS05585 / Atu1131); $Spec^R$ | This study |
| pBCH181 | pRA301::$P_{chvI}$-*lacZ* (ATU_RS00165 / Atu0034); $Spec^R$ | This study |
| pBCH172 | pET-28a(+)::*chvI* (ATU_RS00165 / Atu0034); N-terminal His tag on ChvI; $Km^R$ | This study |
| pBCH185 | pET-28a(+)::ChvID52E (ATU_RS00165 / Atu0034);)); N-terminal His tag on ChvID52E allele; $Km^R$ | This study |
| pBCH175 | pTYB12::*rem* (ATU_RS02820 / Atu0573); N-terminal Intein tag on Rem; $Amp^R$ | This study |
| pPM102 | *exoR* deletion plasmid; pKNG101 derivative, $Strep^R$ | (Tomlinson *et al.*, 2010) |
| pBCH143 | ChvID52N allelic replacement plasmid; pNPTS138 derivative, $Km^R$, $Suc^S$ | This study |
| pBCH144 | ChvID52E allelic replacement plasmid; pNPTS138 derivative, $Km^R$, $Suc^S$ | This study |
| pJX143 | *rem* deletion plasmid; pNPTS138 derivative, $Km^R$, $Suc^S$ | This study |
| pBCH260 | BCH132 allelic replacement plasmid; pNPTS138 derivative, $Km^R$, $Suc^S$ | This study |
| pBCH261 | BCH133 allelic replacement plasmid; pNPTS138 derivative, $Km^R$, $Suc^S$ | This study |
| pBCH262 | BCH134 allelic replacement plasmid; pNPTS138 derivative, $Km^R$, $Suc^S$ | This study |
| pBCH267 | pRA301::$P_{mirA}$ (ATU_RS08050)-*lacZ*; $Spec^R$ | This study |
| pMAT14 | pSRKGm::$P_{mirA}$::*mirA,* (ATU_RS08050), $Gm^R$ | This study |
| pMAT17 | pNPTS::*mirA* deletion fragment allelic replacement plasmid, (ATU_RS08050), $Km^R$, $Suc^S$ | This study |
| pMAT25 | pYW15::*mirA,*(ATU_RS08050), $Amp^R$ | This study |
| pMAT26 | pNPTS::$1638_{S20STOP}$ allelic replacement plasmid, $Km^R$, $Suc^S$ | This study |
| pMAT29 | pNPTS:: $mirA_{G20STOP}$ allelic replacement plasmid, (ATU_RS08050), $Km^R$, $Suc^S$ | This study |
| pMAT37 | pSRKGm:: $P_{mirA}$::$mirA_{S. meliloti;}$ $Gm^R$ | This study |
| pRU2800 | p-$rem_{Sm\ Ru11/001}$; pBBRMCS-5 derivative, cloned into HinDII/BamHI sites, $Gm^R$ | Gift from B. Scharf |
| pMAT39 | pET-28a(+)::*mirA*::$His_6$, (ATU_RS08050), $Km^R$ | This study |

| | | |
|---|---|---|
| pMAT42 | p*mirA* :T25, (ATU_RS08050), Amp$^R$ | This study |
| pMAT43 | p*mirA*::T18, (ATU_RS08050), Km$^R$ | This study |
| pMAT44 | p*rem*::T25 (ATU_RS02820 / Atu0573) | This study |
| pMAT45 | p*rem*::T18, (ATU_RS02820 / Atu0573) Amp$^R$ | This study |
| pMAT46 | p*rem*$_{\text{Receiver domain}}$::T25, (ATU_RS02820 / Atu0573), Amp$^R$ | This study |
| pMAT47 | p*rem*$_{\text{Receiver domain}}$::T18, (ATU_RS02820 / Atu0573), Km$^R$ | This study |
| pMAT48 | p*rem*$_{\text{DNA-binding domain}}$::T25, (ATU_RS02820 / Atu0573), Amp$^R$ | This study |
| pMAT49 | p*rem*$_{\text{DNA-binding domain}}$::T18, (ATU_RS02820 / Atu0573) , Km$^R$ | This study |
| pMAT50 | p*mirA*$_{S.\ meliloti\ \text{1021}}$::T25, (SM_RS17530 / Smb20083), Amp$^R$ | This study |
| pMAT51 | p*mirA*$_{S.\ meliloti\ \text{1021}}$::T18, Smb20083), Km$^R$ | This study |
| pMAT52 | p*rem*$_{S.\ meliloti\ \text{1021}}$::T25, (SM_RS03450, Smc03046), Amp$^R$ | This study |
| pMAT53 | p*rem*$_{S.\ meliloti\ \text{1021}}$::T18, (SM_RS03450, Smc03046), Km$^R$ | This study |
| pMAT54 | pNPTS::*mirA*::FLAG$_3$ allelic replacement plasmid, (ATU_RS08050), Km$^R$, Suc$^S$ | This study |
| pJAS8 | pSRKGm::*remC143A*, (ATU_RS02820 / Atu0573), Gm$^R$ | This study |
| pJAS9 | pSRKGm::*remC143D*, (ATU_RS02820 / Atu0573), Gm$^R$ | This study |
| pJAS10 | pSRKGm::*remC143S*,(ATU_RS02820 / Atu0573), Gm$^R$ | This study |

**Table S4. Oligonucleotides used in this study**

| Primer Name | Sequence 5’ à 3’ | Application |
|---|---|---|
| ChvI D52N Fwd | *CAGCTTGCCATTTTC**AAC**ATTAAGATGCCCCGC* | ChvID52N allele cloning |
| ChvI D52N Rev | *GCGGGGCATCTTAAT**GTT**GAAAATGGCAAGCTG* | |
| ChvI D52E Fwd | *CAGCTTGCCATTTTC**GAG**ATTAAGATGCCCCGC* | ChvID52E allele cloning |
| ChvI D52E Rev | *GCGGGGCATCTTAAT**CTC**GAAAATGGCAAGCTG* | |
| ChvI replace 5' SpeI | actagtATGCAGACGATCGCGCTTGTC | ChvI allele replacement |
| ChvI replace 3' SphI | gcatgcTTAAGCCGCTTCGCGGAA | |
| ChvI seq 1 | AAATCCAGCCGCAGCGTTTT | Verifying site-directed mutation |
| ChvI rev seq 6 | CGAAGACGCTTGATGTGGCT | |
| ChvI comp. 5’ NdeI | catatgCAGACGATCGCGCTTGTCGAC (Heckel, 2014 #681) | Cloning into expression vectors & verifying alleles |
| ChvI comp. 3’ KpnI | ggtaccCTGCTTGAGTCGGCTTC (Heckel, 2014 #681) | |
| ChvI 3' protein exp. XhoI | ctcgagTTAAGCCGCTTCGCGGAAGCGATA | |
| Rem D45N Fwd | *GCGGCAGACAGC**AAC**CTGGCGGCCGTGGAG* | RemD45N allele cloning |
| Rem D45N Rev | *CTCCACGGCCGCCAG**GTT**GCTGTCTGCCGC* | |
| Rem E50N Fwd | *GATCTGGCGGCCGTG**AAC**GCCTTCCTGATC* | RemE50N allele cloning |
| Rem E50N Rev | *GATCAGGAAGGC**GTT**CACGGCCGCCAGATC* | |
| Com-*rem*-P1 NdeI | catatgATCGTAGTGGTTGATGAGCGC (Xu, 2013 #1883) | Rem protein expression |
| Rem 3' protein exp. XhoI | ctcgagTCATTGCCAATCAATGCTGTAGCC | |
| $P_{chvI}$ Fwd KpnI | ggtaccCATGACGCGTTCCTTGTAAAT | *Pchv*I-*lacZ* reporter |
| $P_{chvI}$ Rev SphI | gcatgcCATGTAGTTGGTCTCCATCGT | |
| $P_{aopB}$ Fwd KpnI | ggtaccCATACACGCCCTATTTGCTGG | *PaopB*-*lacZ* reporter |
| $P_{aopB}$ Rev KpnI | ggtaccCATGTTATTCTCCTTTCAGGA | |
| $P_{mirA}$ Fwd KpnI | GGTACCGATTATCAGCGTTGCCGCAGC | |
| $P_{mirA}$ Rev SphI | GCATGCCATCCTTTGGTCTCC CTGTTTTCC | |
| $P_{hcp}$ 5’ entire KpnI | ggtaccCAAGAGTAGTCTATCCCCAG | *Phcp*-lacZ reporters and EMSA amplification |
| $P_{hcp}$ 3’ entire KpnI | ggtaccCATGCAATGAGCCTCCTG | |
| $P_{hcp}$ trunc 116 KpnI | TAAATTGTGGCGTCTACT | |
| $P_{hcp}$ trunc 127 KpnI | GTCTACTCCCTAATCTAA | |
| $P_{hcp}$ trunc 132 | GTGTCTAAATTGTGGCGTCTACTC | *Phcp* EMSA amplification |
| $P_{hcp}$ trunc 140 | TTCAAAATGTGTCTAAATTGTGGC | |
| $P_{hcp}$ trunc 165 | TGAGGCTATTAATCAGACAGACTT | |
| $P_{hcp}$ trunc 195 | ATGGATTTTTTGTAGTTACTTCCC | |

| | | |
|---|---|---|
| pRA301 LB | TGGTGTAAACAAATTGACGCT | |
| $P_{motA}$ P1 HindIII | aagcttCATTCTCACAGTGAAGTTACTACCGC | *PmotA* EMSA amplification |
| $P_{motA}$ P2 EcoRI | gaattcTGAAATCGTCGATGGATTTCACGC | |
| $P_{rem}$ P1 EcoRI | gaattcGCAACCTGAAATTGTCTTTC | *Prem* EMSA amplification |
| $P_{rem}$ P2 HindIII | aagcttCATTCGTCCGCCTCCGAATC | |
| $P_{flgB}$ 5’ | CTGACAATGCCTTGCTGACGAAGA | PflgB EMSA amplification |
| $P_{flgB}$ 3' | CATAGACTTCTCCATCTCGTATGG | |
| *flgB* RT 5’ Phos 5’ RACE | 5PhosTCGAAGGGACTGACA | TSS mapping of *flgB* |
| *flgB* A1 5'RACE | GCTCAGCCATTCGGCTTG | |
| *flgB* A2 5'RACE | TCGAACAGTTGAATCGGTTG | |
| *flgB* S1 5'RACE | GGAAGTCGTGGCGACCAA | |
| *flgB* S2 5'RACE | AACGCCAACACACCCAAGTT | |
| *motA* RT 5'Phos 5'RACE | 5PhosCCATGATGAAGCCGC | TSS mapping of *motA* |
| *motA* A1 5'RACE | ATGATGCAGCCGAAGGTGAT | |
| *motA* A2 5' RACE | ATTCATTGATCGACCACGCC | |
| *motA* S1 5'RACE | ATGGGCGGTCATTTGAACGT | |
| *motA* S2 5'RACE | CGAATTGATGATCATCGGAGGC | |
| *chvI* RT 5'Phos 5'RACE | 5PhosAAATGGCAAGCTGCG | TSS mapping of *chvI* |
| *chvI* A1 5'RACE | CCGAAGTGAGAATATTCCGG | |
| *chvI* A2 5'RACE | GATCGTCTGCATGTAGTTGG | |
| *chvI* S1 5'RACE | AAGGCTACCGGGTCGAAAC | |
| *chvI* S2 5'RACE | TTGACGGGTTGATCGCC | |
| *aopB* RT 5'Phos 5'RACE | 5PhosGGACCAATCCTTGAC | TSS mapping of *aopB* |
| *aopB* A1 5'RACE | TAAGCAGCCGAAAAACCG | |
| *aopB* A2 5'RACE | GGGTTGCTACGAAAATACGC | |
| *aopB* S1 5'RACE | CGACGCCGTAAATGAGGT | |
| *aopB* S2 5'RACE | ACCGGTAGCCTACGACCA | |
| *hcp* RT Primer - 5' RACE | GATACCGGCATCTAC | TSS mapping of *hcp* |
| *hcp* A1 - 5' RACE | AGGTCTGGCTCAAGCGCTCC | |
| *hcp* A2 - 5' RACE | GCGATTGAGATCGATATGCGACAT | |
| *hcp* S1 - 5' RACE | TCGAAGCTGCGGCTTCCGTT | |
| *hcp* S2 - 5' RACE | GGGGCACAGGTACGTGGATA | |
| BCH suppress 5' replace SpeI | actagtTGCGTGAGATGATCCGCA | Suppressor SNP replacement |
| BCH suppress 3' replace SphI | gcatgcCTGCCAGTGATATGCGGTGCT | |
| *bolA* delete P1 SpeI | actagtCAACGAGACCAATGGTCAGGCGAT | *bolA* deletion |
| *bolA* delete P2 | *aagcttggtaccgaattc*CATGGTGTTTCTCCTTAATTCATT | |

| | | |
|---|---|---|
| *bolA* delete P3 | *gaattcggtaccaagctt*TGATTGTGGCCGTGAACG TGCCGG | |
| *bolA* delete P4 PstI | ctgcagTGAGGTTCTGGTGGTGTCGCACAG | |
| pSRK Rev | GTCAATTATTACCTCCACGGGGA | Constructing and confirming cloned constructs |
| pSRK Fwd | GCGTTGGCCGATTCATTAATGCA | |
| pUC/M13 Fwd | CGCCAGGGTTTTCCCAGTCACGAC | |
| pUC/M13 Rev | TCACACAGGAAACAGCTATGAC | |
| MT49 SpeI | ACT AGT CAG CGT TGC CGC | Deletion of *mirA* |
| MT50 | AAGCTTGGTACCGAATTC CCTTTGGTCTCC | |
| MT51 | GAATTCGGTACCAAGCTT TCC CCT TCT GAT | |
| MT52 SphI | GCATGC CTGCTTCGCGAG | |
| MT76 NdeI | CATATG GGC TGA CGG AAG GCT CTG AAA AAG AGC | $P_{mirA}$-*mirA* expresision |
| *mirA* comp 3' SpeI | actagtGGCATCAGAAGGGGATCAGGA | |
| MT135 KpnI | GCGGTACCATGCGGGCCGCAATTAATTC | $P_{N25}$-*mirA* expression |
| MT136 HindIII | GCAAGCTTCATCAGAAGGGGATCAGG | |
| MT177 | CTGGTGCCGCGCGGCAGCCACATATGCTA ATAACGAAGGAGATATACCATGC | pET28a::*mirA*-$His_6$ expression |
| MT178 | AGTGGTGGTGGTGGTGGTGCCTCGAGGAG GAGAGGGGG | |
| MT179 | ACGCCAAGCTACGTAATACGACTCACCATG GACTATCAGGCCG | Allelic replacement construct for Atu1638$_{S1920STOP}$ |
| MT180 | TCACTTTATGTTCTAGAGCATTATTGGTGGC | |
| MT181 | GCCACCAATAATGCTCTAGAACATAAAGTG AACATTATGGAGG | |
| MT182 | GCCTTGACTAGAGGGTCGACGCATGCGTC GACCGTGAATTCAAAACC | |
| MT185 | GCCAAGCTACGTAATACGACTCAGAGATTA CTCATCAACATATCGCTTTTCATTCTC | Allelic replacement construct for *mirA*$_{G2020STOP}$ |
| MT186 | GAATTGGCACTTCTACTCGGCCTGATAGTC C | |
| MT187 | GGACTATCAGGCCGAGTAGAAGTGCCAATT CGCAC | |
| MT188 | CTTGACTAGAGGGTCGACGCATGGGGAGC GGGCGAC | |
| MT224 | TCGACTCTAGAGGATCCCCGGGTAcgggccgc aattaattcccc | pKNT25 and pUT18 sequencing primers |
| MT228 | gcgggcagtgagcgcaacgc | |
| MT229 | CCTTCGCCACGGCCTTGATG | |
| MT230 | CGCCTCGGTGCCCACTGC | MirA-T25 and T18 |
| MT225 | TGATGCGATTGCTGCATGGTCATTGGggaga gggggcgttgg | |
| MT227 | CCGCCCGTGGCCTCGCTGGCGGCggagagg gggcgttgg | |

| | | |
|---|---|---|
| MT235 | TCGACTCTAGAGGATCCCCGGGTAatcgtagtggttgatgagcgcgag | Rem-T25 and T18 |
| MT236 | TGATGCGATTGCTGCATGGTCATttgccaatcaatgctgtagccgaggaaccg | |
| MT250 | CCGCCCGTGGCCTCGCTGGCGGCttgccaatcaatgctgtagccgag | |
| MT255 | TCGACTCTAGAGGATCCCCGGGTAatcgtagtggttgatgagcgc | Rem receiver domain-T25 and T18 |
| MT259 | TGATGCGATTGCTGCATGGTCATgcggatggccgc | |
| MT256 | CCGCCCGTGGCCTCGCTGGCGGGgcggatggccgc | |
| MT257 | TCGACTCTAGAGGATCCCCGGGTAtcgaacttcaccgatgcag | Rem DNA-binding domain-T25 and T18 |
| MT260 | TGATGCGATTGCTGCATGGTCATccaatcaatgctgtagccgagg | |
| MT258 | CCGCCCGTGGCCTCGCTGGCGGCccaatcaatgctgtagccgag | |
| MT284 | TCGACTCTAGAGGATCCCCGGGTAatcgtggtggttgatgatagagc | Rem (*S. meliloti* 1021)-T25 and T18 |
| MT285 | TGATGCGATTGCTGCATGGTCATattccagtcgatgcagtagcc | |
| MT287 | CCGCCCGTGGCCTCGCTGGCGGCattccagtcgatgcagtagcc | |
| MT288 | TCGACTCTAGAGGATCCCCGGGTACCTGTCATCAGCCCCACG | MirA (*S. meliloti* 1021)-T25 and T18 |
| MT289 | TGATGCGATTGCTGCATGGTCATGTGCAAGCTCTTGGAGTGACTGCGCAG | |
| MT291 | CCGCCCGTGGCCTCGCTGGCGGCGTGCAAGCTCTTGGAGTGACTG | |
| MT279 | atgTCGGAAAACAGGGAGACCAAAGgATGCCTGTCATCAGCCCCACGATCC | Construction of P*mirA*$_{A. tumefaciens}$- MirA (*S. meliloti* 1021) Expression vector |
| MT264 | cgggggatccactagTCAGTGCAAGCTCTTGGAGTG | |
| MT261 | ctagtggatcccccgggctgc | |
| MT262 | cctttggtctcccttggacgttaacggcaagtctcg | |

| | | |
|---|---|---|
| Rem_C143A_Q5_1 | TCCTGAAGTTgccGGCGAAGTCTTTG | Construction of *remC143A* |
| Rem_C143A_Q5_2 | TCGCGACCATCGGAGAAA | |
| Rem_C143D_Q5_1 | TCCTGAAGTTgacGGCGAAGTCTTTG | Construction of *remC143D* |
| Rem_C143D_Q5_2 | TCGCGACCATCGGAGAAA | |
| Rem_C143S_Q5_1 | TCCTGAAGTTagcGGCGAAGTCT | Construction of *remC143S* |
| Rem_C143S_Q5_2 | TCGCGACCATCGGAGAAAAC | |

**Supplementary Figures Legends**

**Figure S1. Alignment of Rem, ChvI, and other response regulator proteins.** ClustalW (Clustal 2.1) was used to align the amino acid sequence of Rem, ChvI, and other response regulator proteins. The predicted receiver domain is highlighted in light gray and DNA-binding domain is highlighted in dark gray. The conserved aspartate that receives the phosphate during signal transduction is labeled in red; for ChvI this is D52, for Rem proteins, this position contains a glutamic acid residue E50. The D45 residue for Rem proteins is labeled in yellow. An asterisk indicates residues of conserved identity; a colon indicates conservation of residues with strongly similar properties; a period indicates conservation of residues with weakly similar properties. Amino acid sequences of proteins can be found at NCBI reference numbers WP_006313038.1 (Rem/*A. tumefaciens*), CAC45250.1 (Rem/*S. meliloti*), SUB44985.1 (Rem/*O. anthropi*), WP_006698188.1 (Rem/*R. pusense* IRGB74), WP_065114859.1 (Rem/*A. rhizogenes*) WP_010970615.1 (ChvI/*A. tumefaciens*), WP_003531999.1 (ChvI/*S. meliloti*), AYM61105.1 (VirG/*A. tumefaciens*), NP_417864.1 (OmpR/*E. coli*), and AKH06072.1 (PhoB/*S. enterica*).

**Figure S2. Calcofluor binding by *chvI* phosphorylation site mutants.** *chvID52E* phenocopies Δ*exoR* for Calcofluor binding, while *chvID52N* phenocopies Δ*chvI.* Fluorescence of solid culture bacterial growth from 5 μl of cell suspension ($OD_{600}$=0.2) on ATGN agar supplemented with 200 μg/ml Calcofluor illuminated by UV light after 48 h incubation. Calcofluor binds to the exopolysaccharide succinoglycan, the production of which is elevated in Δ*exoR* (Heckel *et al.*, 2014, Tomlinson *et al.*, 2010).

**Figure S3. Deletion of *exoR* does not impact $P_{visN}$ or $P_{rem}$ promoter activity.** *β*-galactosidase assay of strains grown to exponential phase, followed by lysis and measurement of promoter activity. Bars represent the mean of three replicates, with error bars representing standard deviation.

**Figure S4. ChvI-dependent promoter mapping and ChvI binding site discovery**. Electrophoretic mobility shift assays to determine the minimum DNA sequence required for $His_6$-ChvI$^{D52E}$ binding to the T6SS intergenic region. Fragment lengths relative to the *clpV* start codon are indicated. 10 nM DNA was incubated with or without 600 nM purified $His_6$-ChvI$^{D52E}$ and binding was assessed by migration of DNA when run on a 6% polyacrylamide gel. The gel was post-stained with SYBR Safe dye for DNA detection.

**Figure S5. Rem activity is phosphorylation-independent.** (A) Motility of strains harboring *rem* alleles under control of the *lac* promoter was assessed by inoculating colonies into 0.3% agar -/+ 400 µM IPTG and measuring swim ring diameter, three days post-inoculation. Presented relative to the uninduced wild-type strain carrying each expression construct. Error bars represent standard deviation for at least three replicates. (B) Biofilm forming capabilities of the same strains tested for motility were assessed. Strains were inoculated -/+ 400 µM IPTG and grown 48 h before measuring biofilm biomass. Biofilm biomass was calculated by $A_{600}$ absorbance of 0.1% crystal violet-dyed biofilms solubilized in 33% acetic acid, divided by the $OD_{600}$ of each culture. Data are presented as the biofilm biomass relative to uninduced wild-type C58 carrying each allele construct. Error bars represent standard deviation of the mean for at least three replicates.

**Figure S6. A single cysteine residue in Rem is not required for motility.** Motility of strains harboring *rem* alleles under control of the $P_{lac}$ promoter was assessed by inoculating colonies into 0.3% agar -/+ 500 µM IPTG and measuring swim ring diameter after 48 h incubation in a humid chamber at RT. Error bars represent standard deviation for at least three replicates.

**Figure S7. Transcription start sites for *motA* and *flgB* were mapped by 5' RACE**. (A) Alignment of five Rem-regulated (class II) flagellar gene promoters and identification of a putative Rem binding site. Promoters were chosen based on Rem-regulated promoters in *S. meliloti* (Rotter *et al.*, 2006). The relative positions of the transcription and translation start sites are indicated. Conserved positions are highlighted (Dark gray, 100% conservation, light gray, >60% conservation). Direct repeats are indicated by arrows. Transcription start sites of $P_{flgB}$ and $P_{motA}$ are labeled in green; sites were mapped by 5' RACE. (B) A 262 bp region upstream of the *flgB* (Atu0555) gene was assayed for Rem binding. Promoter DNA for the ChvI-regulated gene *aopB* was used as a negative control. A single shift is observed (indicated by asterisk) for Rem binding to $P_{flgB}$. C. Purified Rem incubated with *rem* promoter DNA (283 bp upstream of Rem start codon) does not produce a discrete shift when assayed with the same conditions for A. Incubation with $P_{motA}$ DNA was used as a positive control (shifts indicated by asterisks).

**Figure S8. ChvI does not interact with Rem or alter Rem-DNA binding.** (A) $His_6$-ChvI$^{D52E}$ does not disrupt Rem-$P_{motA}$ interactions. Rem was incubated with $P_{motA}$ 10 min prior to addition of $His_6$-ChvI$^{D52E}$ DNA, followed by 20 min incubation prior to electrophoretic separation on a 6% acrylamide gel and DNA-staining with SYBR safe dye. (B) Western (left) and farwestern (right) blot analysis for $His_6$-ChvI$^{D52E}$-Rem interaction on a nitrocellulose membrane. $His_6$-ChvI$^{D52E}$ (positive control), Rem, and BSA (negative control) were separated on a polyacrylamide gel and transferred to a nitrocellulose membrane. For the western blot (left), proteins were detected with anti-$His_6$ antibody; for the farwestern (right), immobilized proteins were denatured with guanidine-HCl and renatured prior to probing with $His_6$-ChvI$^{D52E}$, followed by detection with anti-$His_6$ antibody.

**Figure S9.** The **Δ*exoR* and Δ*rem* mutants have distinct biofilm defects, and loss of *exoR* is epistatic to Δ*rem*.** Biofilm formation ($A_{600}/OD_{600}$) of strains was measured after 48 h incubation at RT in a humid chamber by staining biofilm cells with 0.1% crystal violet and solubilization in 33% acetic acid. Error bars represent standard deviation of three replicates.

**Figure S10. Characterization of motile suppressors of Δ*exoR.*** (A) Suppressor mutants BCH132, BCH133, and BCH134 lose motility inhibition by ExoR-ChvG-ChvI but are not rescued for biofilm formation. (B). Activity of translational fusion to $P_{rem}$, $P_{chvI}$, and $P_{aopB}$ in wild-type, Δ*exoR* (Δ*exoA*) strains, and BCH133. (C) Motility of the recreated suppressor mutation allele from BCH133 $mirA_{FS}$ (re-created in naïve background) in wild type (BCH136) and Δ*exoR*(Δ*exoA*) $mirA_{FS}$ (re-created in naïve background) (BCH137). (D) β-galactosidase activity of $P_{mirA}$ in BCH136 and BCH137.

**Figure S11. Mutation of *bolA* does not impact motility or biofilm formation. (**A) Biofilm formation ($A_{600}/OD_{600}$) of Δ*bolA* strain relative to wild-type and Δ*exoR* (Δ*exoA*) strains; biofilms were stained with 0.1% crystal violet followed by solubilization with 33% acetic acid. Error bars represent standard deviation of three replicates. (B) Motility of Δ*bolA* strain relative to wild-type and Δ*bolA* strains. Strains were inoculated in the center of 0.3% Bacto Agar / ATGN and incubated 24 h in a humid chamber at RT. Error bars represent standard deviation of three replicates.

**Figure S12. Motility suppression of Δ*exoR* occurs following mutation of *mirA,* but not Atu1638.** Motility assay for $Atu1638_{S20STOP}$ (A) and $mirA_{G20STOP}$ (B) in Δ*exoR* (Δ*exoA*) strains. Colonies were inoculated to the center of ATGN 0.3% Bacto Agar and incubated for 24 h in a humid chamber at RT. Error bars represent standard deviation from the mean of at least three replicates.

**Figure S13.Ectopic expression of *mirA* leads to motility inhibition.** The *mirA* gene was expressed from the native *mirA* promoter with pMAT14 (pSRKGM::$P_{mirA}$::*mirA*) in wild-type, Δ*exoR* (Δ*exoA*), and Δ*mirA* strains. Colonies were inoculated to the center of ATGN 0.3% Bacto Agar and incubated for 24 h in a humid chamber at RT. Error bars represent standard deviation from the mean of at least three replicates.

**Figure S14. Pre-incubation with MirA does not inhibit His$_6$-ChvI$^{D52E}$ DNA binding.** His$_6$-ChvI$^{D52E}$ was incubated with MirA 15 min at room temperature prior to the addition of 5 nM $P_{hcp}$ DNA, followed by 15 min incubation at RT. Products were electrophoresed on an 8% acrylamide gel (80:1 acrylamide:bisacrylamide). DNA was labeled with SYBR Green dye and visualized by BioRad ChemiDoc.

**Figure S15. Alignment of putative MirA homolog proteins from different members of *Rhizobiales* order.** ClustalW (Clustal 2.1) was used to align the amino acid sequence of proteins. An asterisk indicates residues of conserved identity; a colon indicates conservation of residues with strongly similar properties; a period indicates conservation of residues with weakly similar properties. The amino acid sequences correspond to the following NCBI reference numbers: WP_003502959.1 (MirA/ *A. tumefaciens*), CAC48483.1 (MirA/ *S. meliloti*), WP_004442019.1 (MirA/*R. pusense* IRGB74), WP_065115793.1 (MirA/*A. rhizogenes*), and WP_040128199.1 (MirA/*Ochrobactrum/Brucella anthropi*).

**Figure S16. *A. tumefaciens* and *S. meliloti rem* genes can cross-complement for motility.** Colonies were inoculated to the center of ATGN 0.3% Bacto Agar (for *A. tumefaciens* strains) or Bromfield medium 0.3% Bacto Agar (for *S. meliloti* strains) and incubated 24 h in a humid chamber at RT. Error bars represent standard deviation from the mean of at least three replicates.

#### Rem and ChvI Response Regulator Alignment

```
A_tum_Rem       --------------MIVVVDERELVKDGYTSLFGREGIPSTGFDPVEFGEWVSTAADSDL
S_mel_Rem       --------------MIVVVDDRALVKDGYASLFGREGIPSTGFDPREFGEWVSSAADSDI
O_ant_Rem       --------------MIVVVDDRDIVTEGYSSWFGREGITTTGFTPTDFDEWVESVPQQDI
R_IRB_Rem       --------------MIVVVDERELVKDGYTSLFGREGIPSTGFDPVEFGEWVSTAAESDL
A_rhi_Rem       --------------MIVVVDERELVKDGYTSLFGREGIPSTGFDPAEFGEWVSTAAESDL
A_tum_ChvI      ------------MQTIALVDDDRNILTSVSIALEAEGYRVETYTDGASALDGLIARPPQL
S_mel_ChvI      ------------MQTIALVDDDRNILTSVSIALEAEGYKVETYTDGASALEGLLARPPQL
A_tum_VirG      MAGQDPRLRGEPLKHVLVIDDDVAMRHLIVEYLTIHAFKVTAVADSKQFNRVLCSETVDV
E_col_OmpR      ---------MQENYKILVVDDDMRLRALLERYLTEQGFQVRSVANAEQMDRLLTRESFHL
S_ent_PhoB      -----------MARRILVVEDEAPIREMVCFVLEQNGFQPVEAEDYDSAVNKLNEPWPDL
: ::::   :        :  ..                     .:

A_tum_Rem       AAVEAFLIGQ-GDRSFSLPKAIRDRTTAPVIAVSDQPSLEATLALFDSGVDDVVRKPVHP
S_mel_Rem       DAVEAFLIGQ-GESTFTLPRAIRDRSRAPVIAMSDTPSLENTLALFDCGVDDVVRKPVHP
O_ant_Rem       MAVEAFLIGE-CADQHRLPARIRERCKAPVIAVNDRPSLEHTLELFQSGVDDVVRKPVHV
R_IRB_Rem       AAVEAFLIGQ-GDRSFSLPKAIRDRTTAPVIAVSDQPSLEATLALFDSGVDDVVRKPVHP
A_rhi_Rem       AAVEAFLIGQ-GDRSYSLPKAIRDRTTAPVIAVSDQPSLESTLALFDSGVDDVVRKPVHP
A_tum_ChvI      AIFDIKMPRMDGMELLRRLR---QKSDIPVIFLTSKDEEIDELFGLKMGADDFITKPFSQ
S_mel_ChvI      AIFDIKMPRMDGMELLRRLR---QKSDLPVIFLTSKDEEIDELFGLKMGADDFITKPFSQ
A_tum_VirG      VVVDLNLGREDGLEIVRSLA--TKSDVPIIIISGARLEEADKVIALELGATDFIAKPFGT
E_col_OmpR      MVLDLMLPGEDGLSICRRLR--SQSNPMPIIMVTAKGEEVDRIVGLEIGADDYIPKPFNP
S_ent_PhoB      ILLDWMLPGGSGLQFIKHLKREAMTRDIPVVMLTARGEEEDRVRGLETGADDYITKPFSP
.:  :                       ::      .    :  :. *. * : **.

A_tum_Rem       REILARAAAIRRRLQ-------------VISN----FTDAGPIRVFSDGRDPEVCG-EVF
S_mel_Rem       REILARVAAIRRRLT-------------AIAN----FTDIGPIRVFADGRDPEING-EVF
O_ant_Rem       REILARINAIRRRAG-------------ASAASNDSGTELGPIRVFSDGRDPQING-IDF
R_IRB_Rem       REILARAAAIRRRLQ-------------VISN----FTDAGPIRVFSDGRDPEVGG-EVF
A_rhi_Rem       REILARAAAIRRRLQ-------------VISN----FTDAGPIRVFSDGRDPEVGG-EVF
A_tum_ChvI      RLLVERVKAILRRASSREASAATGGTLKPTADQQARTLERGQLAMDQERHTCTWKG-EPV
S_mel_ChvI      RLLVERVKAILRRAANREAAAGGTGPAKN-ADTPSRSLERGQLVMDQERHTCTWKN-ESV
A_tum_VirG      REFLARIRVALRVRP------------SVARTKDRRSFSFADWTLNLRRRRLISEEGSEV
E_col_OmpR      RELLARIRAVLRRQANE---------LPGAPSQEEAVIAFGKFKLNLGTREMFRED-EPM
S_ent_PhoB      KELVARIKAVMRRIS---------------PMAVEEVIEMQGLSLDPGSHRVMTGD-SPL
: :: *  .  *                                 :    :         .

A_tum_Rem       ALPRRERRILEYLIANRGRRVSKSQIFNAIYGIFDDDVEENVVESHISKLRKKLR-KKLG
S_mel_Rem       ALPRRERRILEYLVANRGRRVSKSQIFNAIYGIFDEDVEENVVESHISKLRKKLR-KKLG
O_ant_Rem       PLPRRERRILEYLIANRGRRLNKAQIFSAIYGIFDSEVEENVVESHISKLRKKLR-EQMG
R_IRB_Rem       PLPRRERRILEYLIANRGRRVSKAQIFNAIYGIFDDDVEENVVESHISKLRKKLR-KKLG
A_rhi_Rem       PLPRRERRILEYLIANRGRRVSKAQIFNAIYGIFDDDVEENVVESHISKLRKKLR-KKLG
A_tum_ChvI      TLTVTEFLILHSLAQRPGVVKSRDALMDAAY-DEQVYVDDRTIDSHIKRLRKKFKLVDGD
S_mel_ChvI      TLTVTEFLILHALAQRPGVVKSRDALMDAAY-DEQVYVDDRTIDSHIKRLRKKFKMVDND
A_tum_VirG      KLTAGEFNLLVAFLEKPRDVLSREQLLIASR-VREEEVYDRSIDVLILRLRRKLEGDPTT
E_col_OmpR      PLTSGEFAVLKALVSHPREPLSRDKLMNLAR-GREYSAMERSIDVQISRLRRMVEEDPAH
S_ent_PhoB      DMGPTEFKLLHFFMTHPERVYSREQLLNHVW-GTNVYVEDRTVDVHIRRLRKALEHSGHD
:   *  :*  :  .      .:  ::       :  . :. ::  * :**: ..

A_tum_Rem       YDPVDSKRFLGYSIDWQ
S_mel_Rem       FDPIDSKRFLGYCIDWN
O_ant_Rem       FDPIDSKRFLGYCINIE
R_IRB_Rem       YDPVDSKRFLGYSIDWQ
A_rhi_Rem       YDPVDSKRFLGYSIDWQ
A_tum_ChvI      FDMIETLYGVGYRFREAA
S_mel_ChvI      FDMIETLYGVGYRFRETA
A_tum_VirG      PQLIKTARGAGYFFDADVDVSYGGVMAA
E_col_OmpR      PRYIQTVWGLGYVFVPDGSKA
S_ent_PhoB      R-MVQTVRGTGYRFSTRF
:.:    **
```

**Alakavuklar et al. FIG S1**

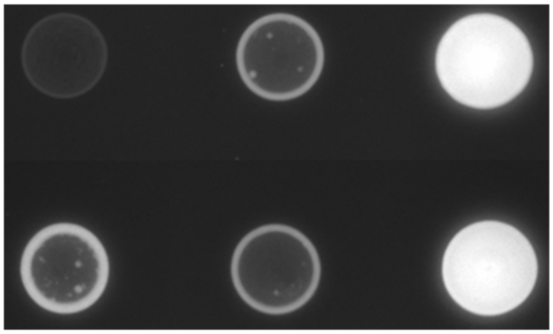

**Alakavuklar et al. FIG S2**

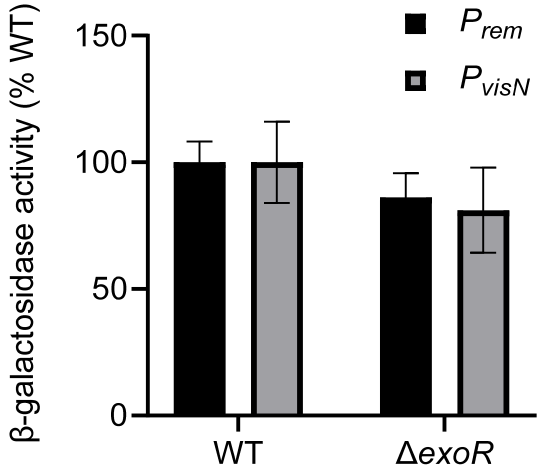

**Alakavuklar et al. FIG S3**

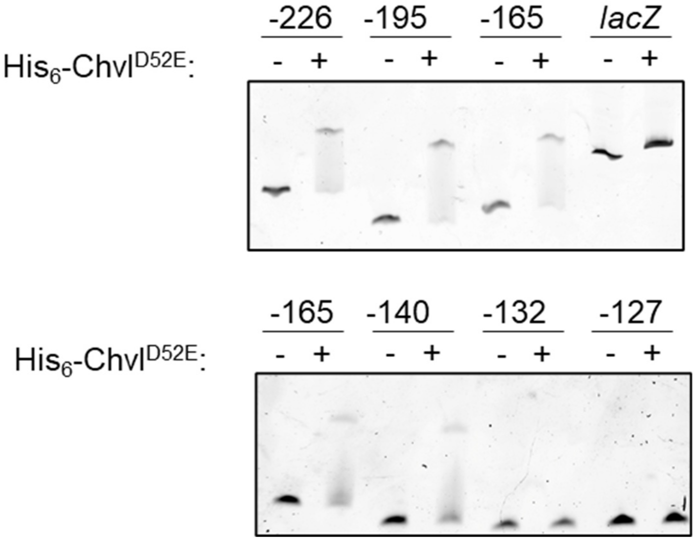

**Alakavuklar et al. FIG S4**

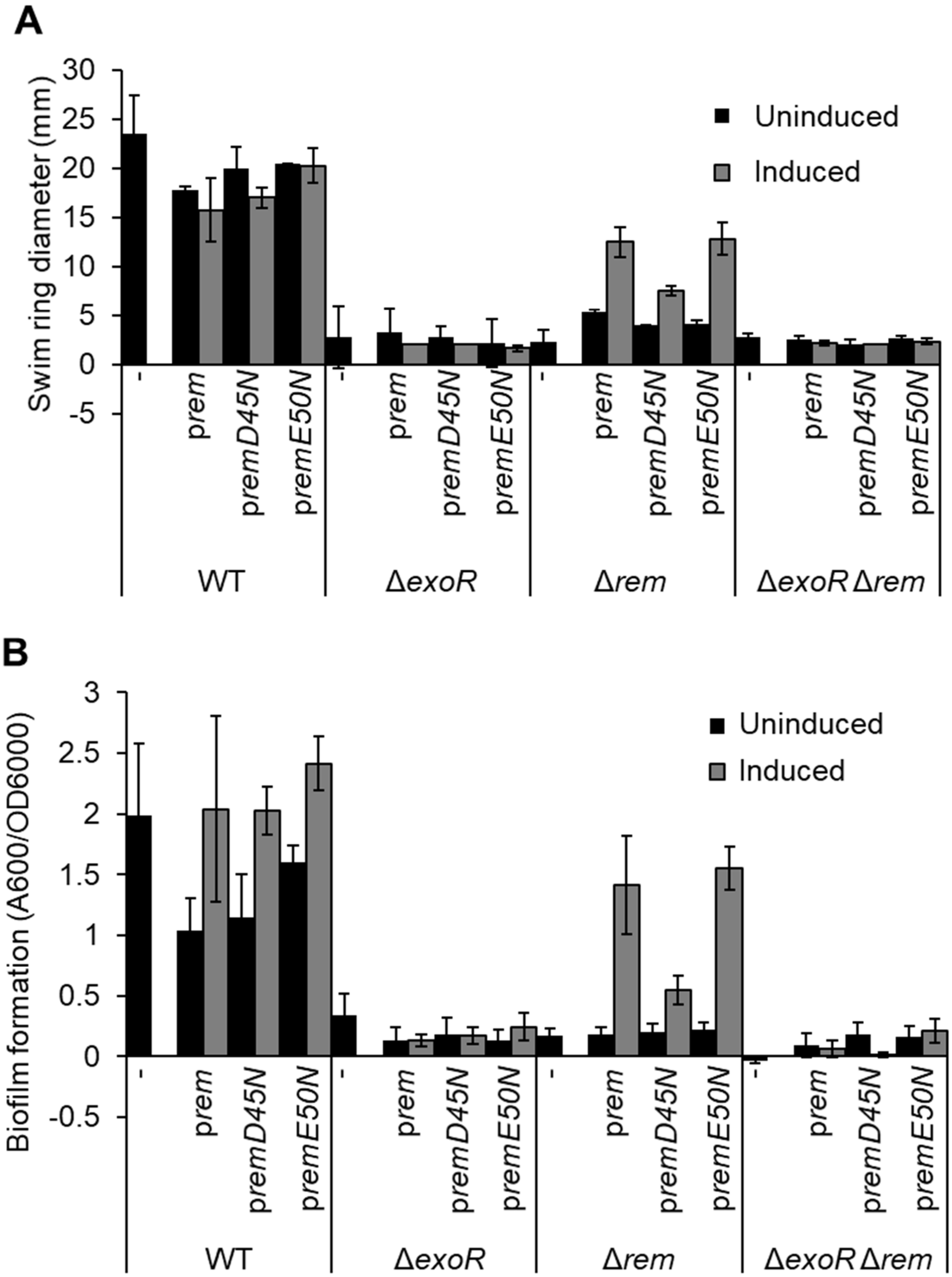

**Alakavuklar et al. FIG S5**

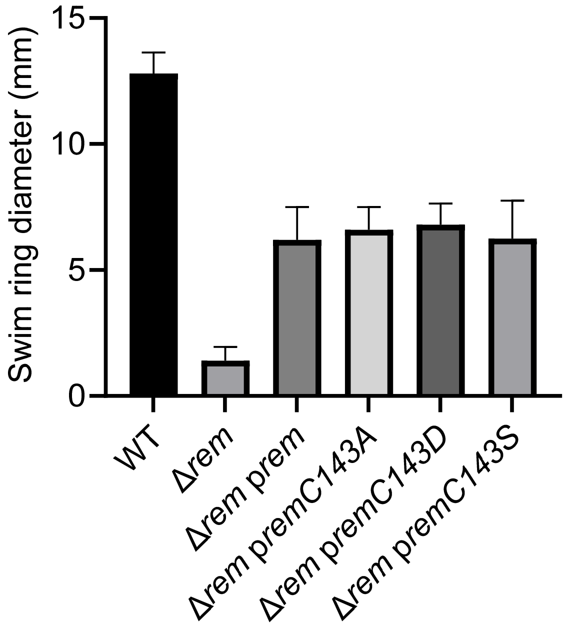

**Alakavuklar et al. FIG S6**

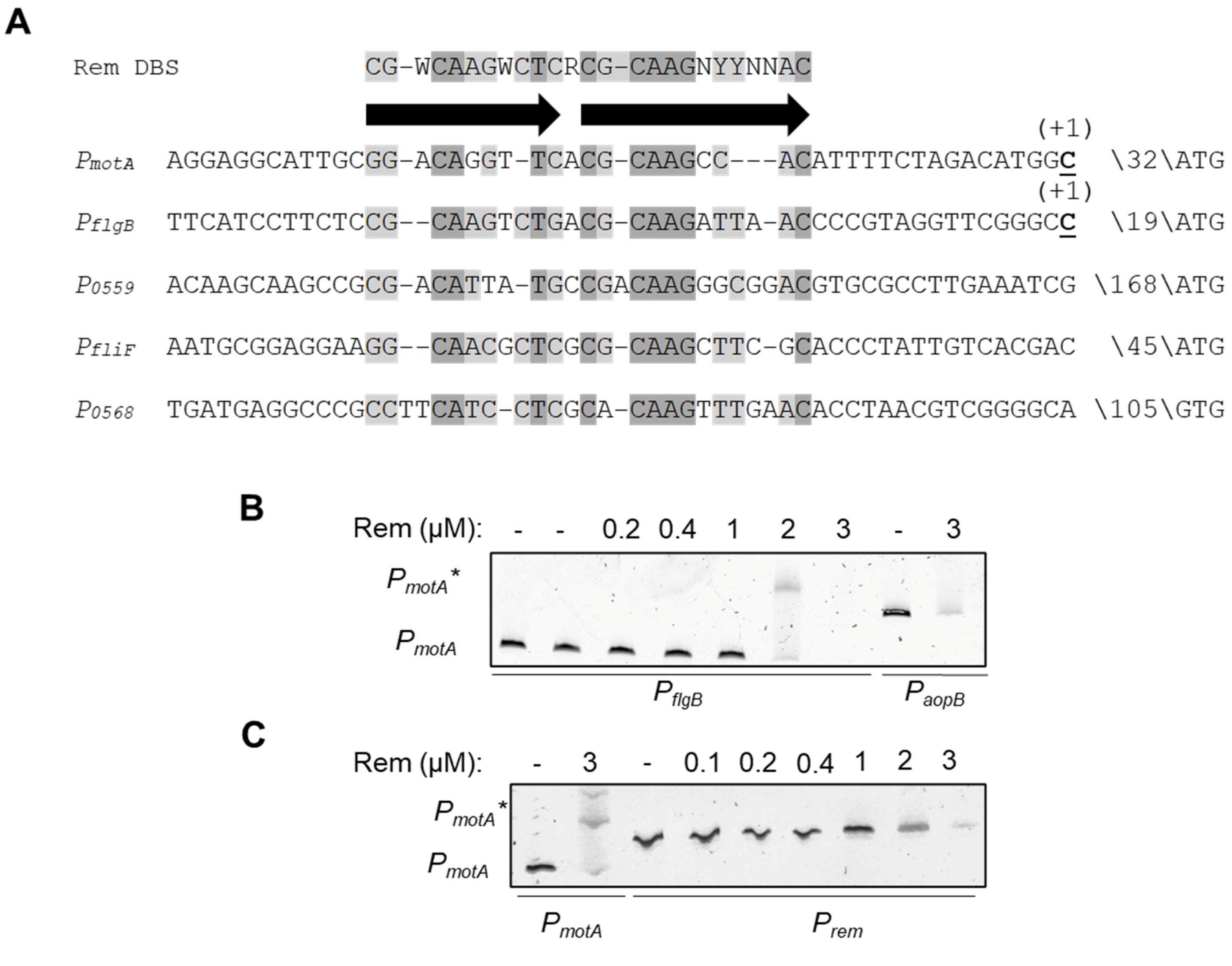

**Alakavuklar et al. FIG S7**

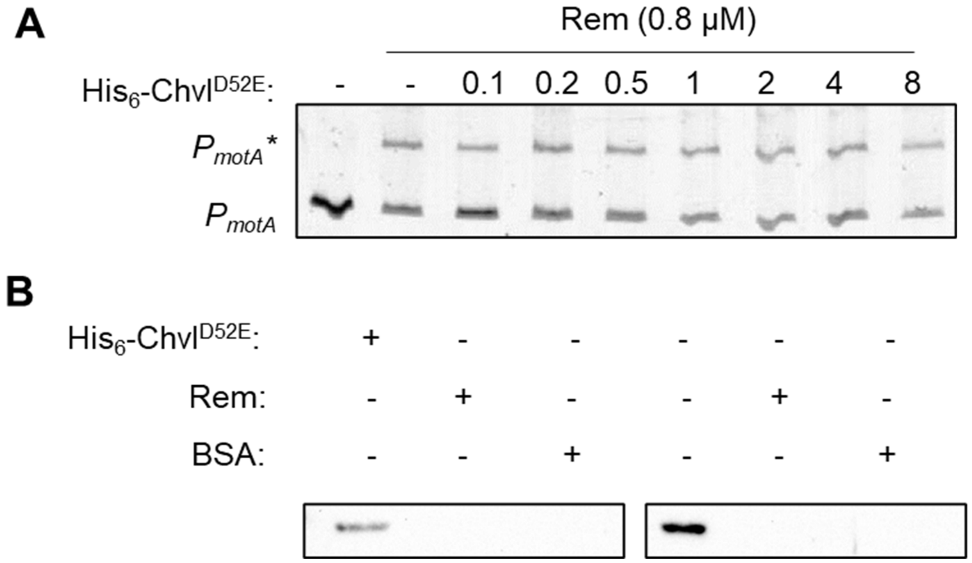

**Alakavuklar et al. FIG S8**

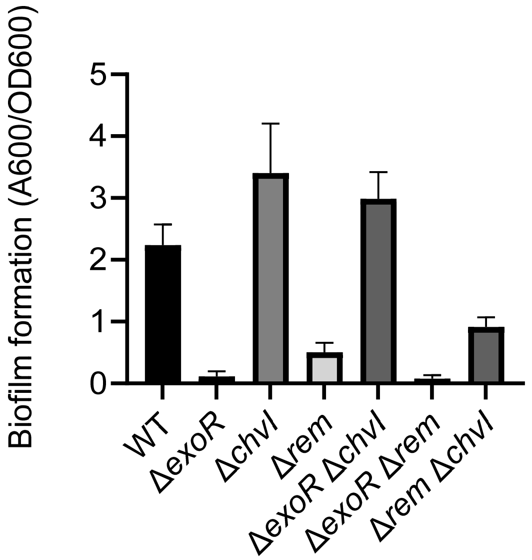

**Alakavuklar et al. FIG S9**

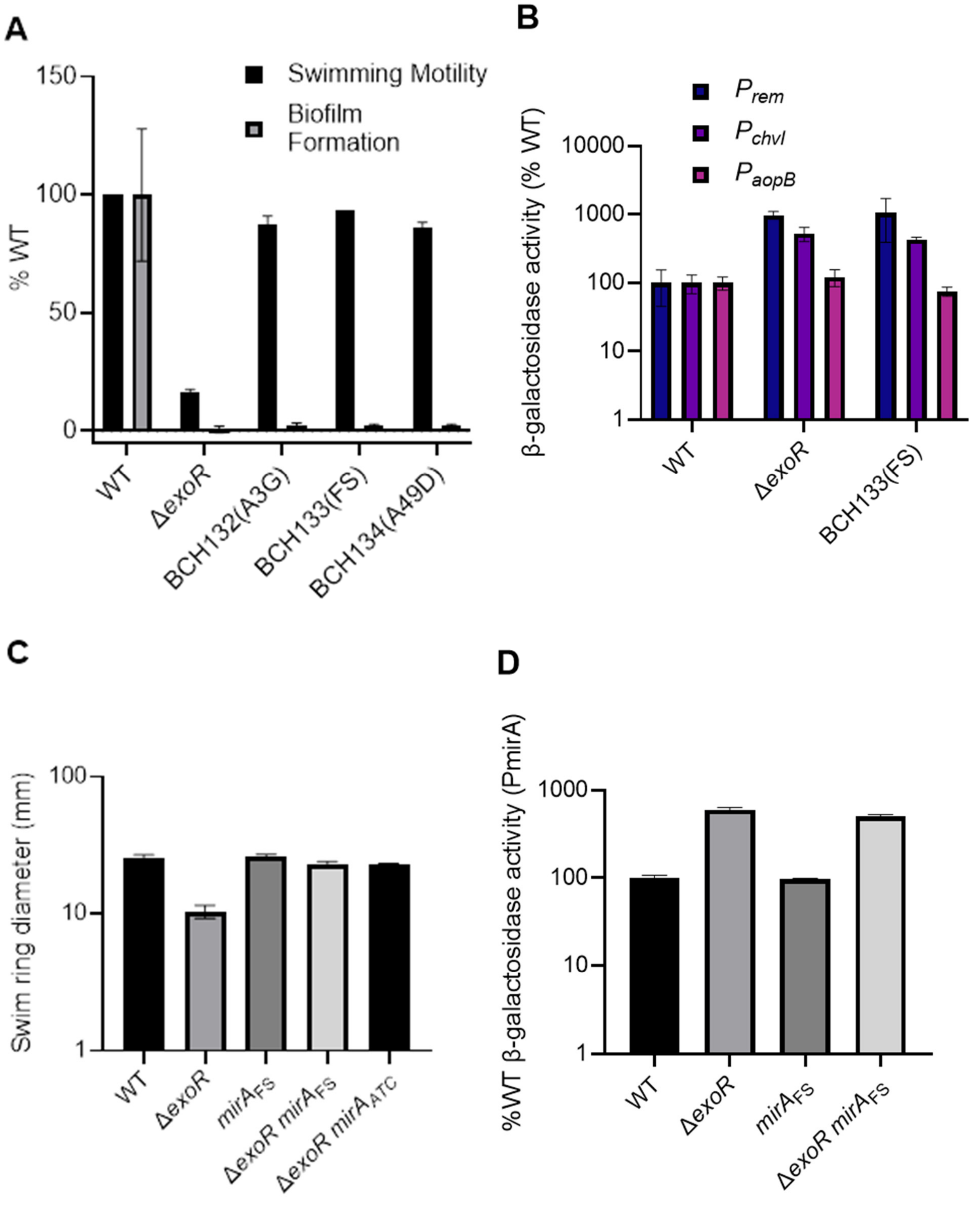

**Alakavuklar et al. FIG S10**

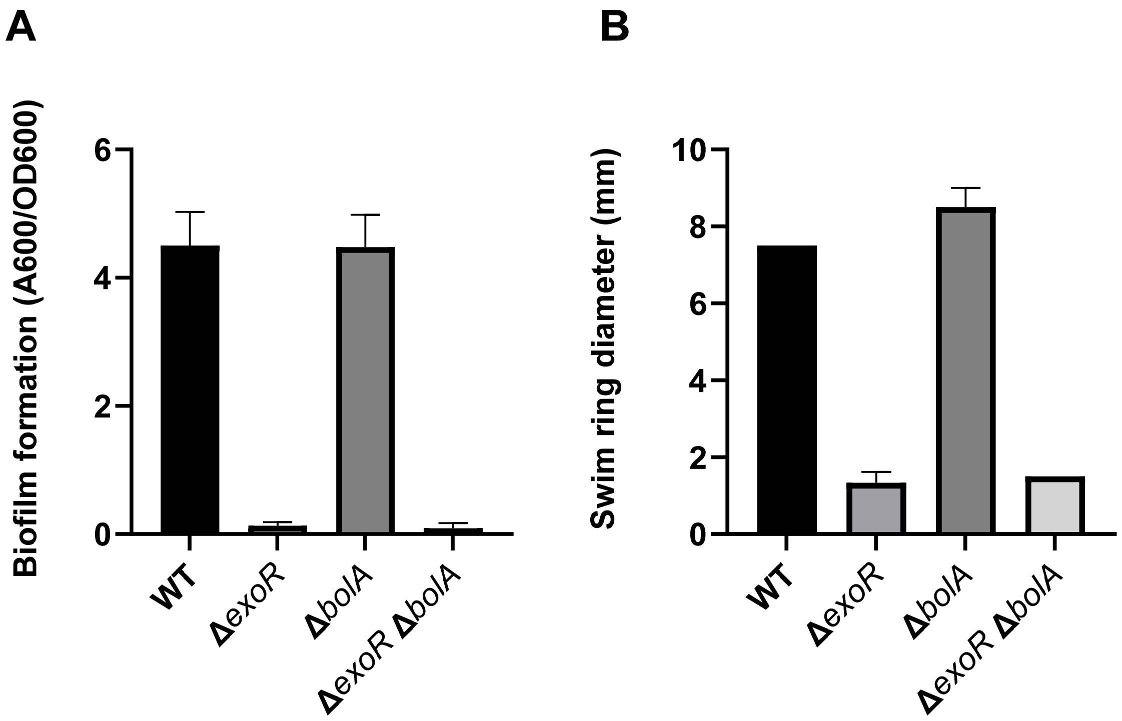

**Alakavuklar et al. FIG S11**

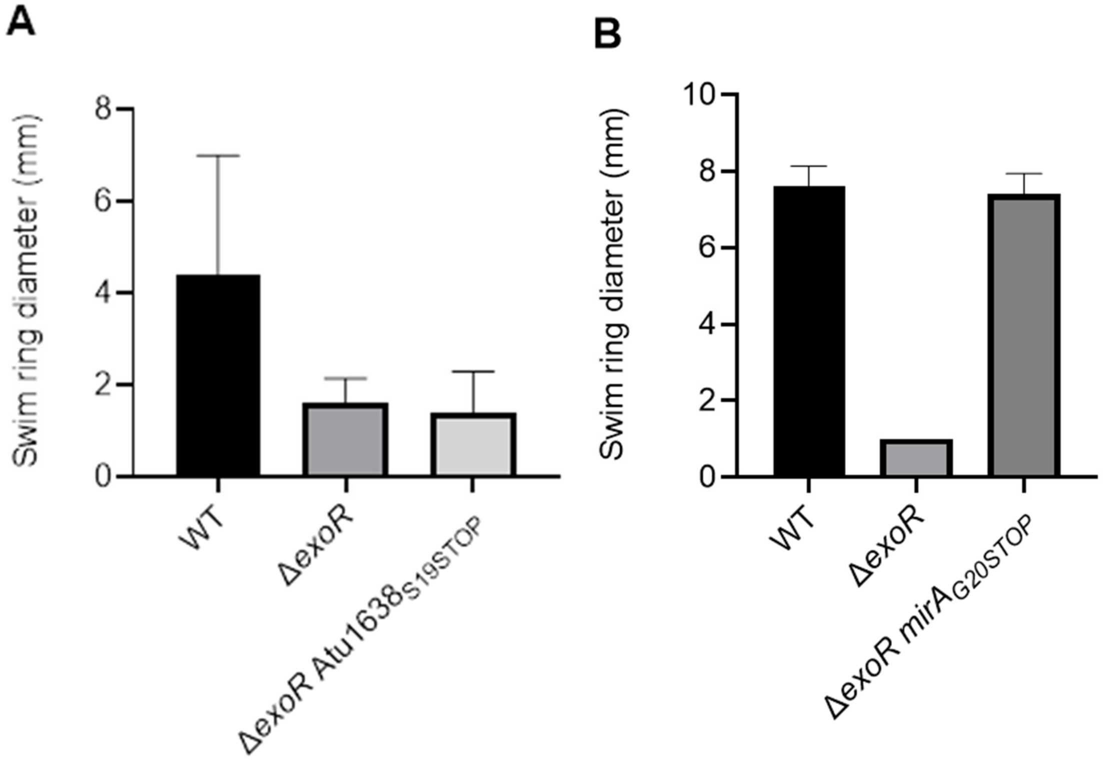

**Alakavuklar et al. FIG S12**

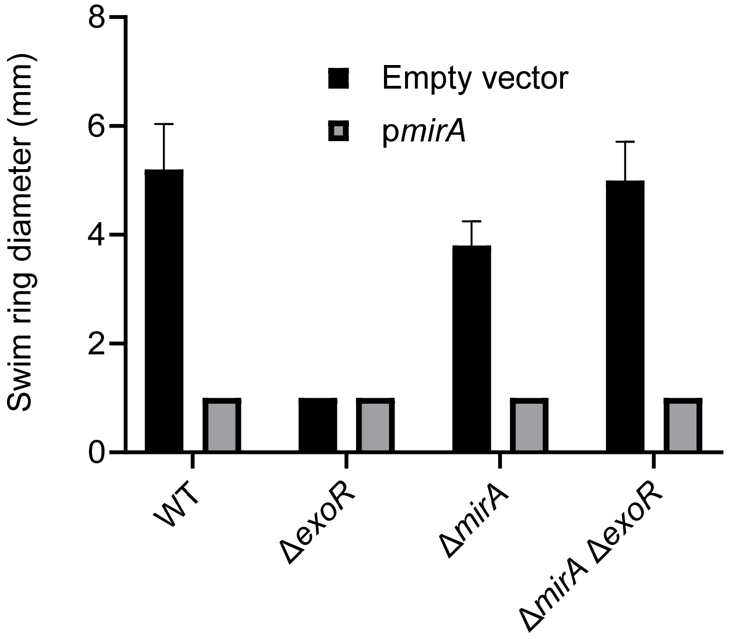

**Alakavuklar et al. FIG S13**

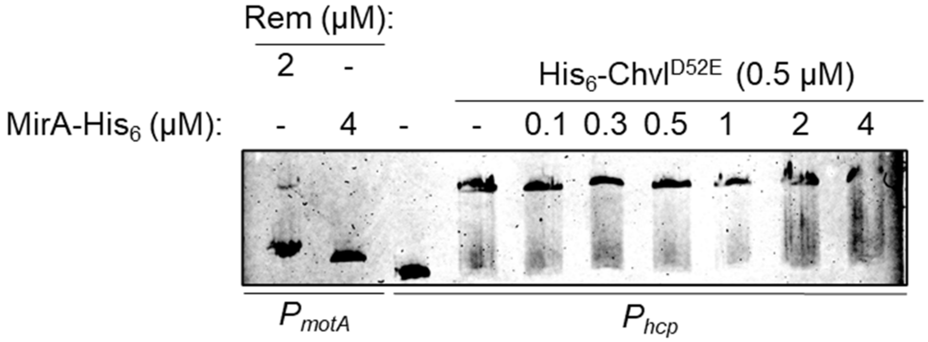

**Alakavuklar et al. FIG S14**

**MirA Protein Homolog Alignment**

```
At_C58  MRAAINSPSLSIDTMDYQAECQFALEPSIQGLIEKAEHAGWNRQQAALAIVALASEHLTDLLTAGGLPAPDQRPLS
R_IRBG  MRAAINSPSLSIDTMDYQAECQFALEPSIQGLIEKAEHAGWNRQQAALAIVALASEHLTDMLST--LAAPDQRPHS
A_rhiz  MRAAINSPSLSIDTMDYQAECQFALEPSIQGLIEKAEDAGWNRQQAALAIVALASEHVTDMLSA--LAAPDQRPLS
O_anth  MGALINPPSKLISAEGDEIECKFALEPSFVCLVSEAKNAGWDELQVAFSLISLCDSIIYGVKPKD
S_meli    MPVISPTIHTQQNSRELECQRAMEDEFLVLTALAEEAGWSWQEIALAMLELTEQYVAAMRPSSAMEARSLLRSHSKSLH
            : .*:   .  . : **: *:* .:  *   *:.***.  : *:::: * .. : : .       .
```

### Alakavuklar et al. FIG S15

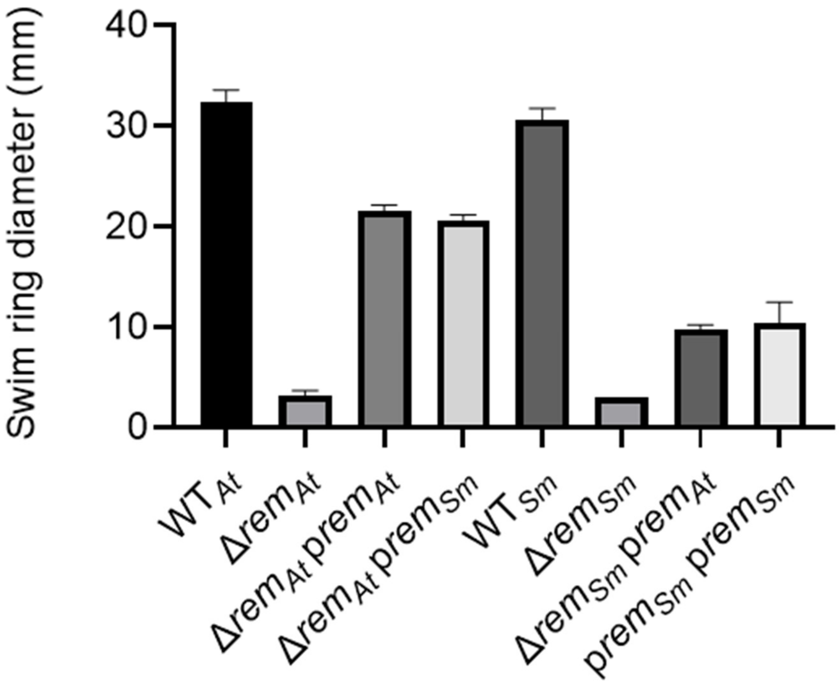

**Alakavuklar et al. FIG S16**
